# Isolation and Characterization of Antibodies Against VCAM-1 Reveals Putative Role for Ig-like Domains 2 and 3 in Cell-to-Cell Interaction

**DOI:** 10.1101/2024.12.03.626733

**Authors:** Binura Perera, Yuao Wu, Jessica R. Pickett, Nadya Panagides, Francisca M. Barretto, Christian Fercher, David P. Sester, Martina L. Jones, Hang T. Ta, Lucía F. Zacchi

## Abstract

Vascular cell adhesion molecule-1 (VCAM-1) plays an important role in inflammation, where it facilitates the recruitment of leukocytes to the inflamed area via leukocytes’ VLA-4 and endothelial cells’ VCAM-1 interaction. VCAM-1 expression is also upregulated in certain cancers. VCAM-1 has 7 Ig-like domains, with domains 1 and 4 shown to be critical for VLA-4 binding. However, the specific functions of individual VCAM-1 Ig-like domains remain poorly understood. In this study, we identified single-chain variable fragment (scFvs) antibodies targeting domains 2, 3, and 5 of VCAM-1, and investigated the ability of these antibodies to block VCAM-1-mediated cell adhesion to macrophages. We show that scFv antibodies against Ig-like domains 2 and 3 significantly interfere with the ability of macrophages to bind endothelial cells, suggesting that these domains also play a role in facilitating this interaction. These results emphasize the need to more carefully study the role of each domain on VCAM-1 function and highlight the potential of targeting these VCAM-1 domains for more tailored therapeutic interventions in inflammatory diseases and cancer.

## 1. Introduction

Vascular cell adhesion protein 1 (VCAM-1) is a cell adhesion molecule implicated in a range of diseases, including atherosclerosis, immunological disorders (e.g. rheumatoid arthritis (RA) and asthma), and cancer, among others [1–3]. VCAM-1 is expressed in a variety of cells and tissues, including endothelial cells. VCAM-1 expression is induced by proinflammatory cytokines [3] and it is involved in the recruitment of leukocytes during inflammation [2]. For example, during atherosclerosis, endothelial cells express VCAM-1, and this assists in the migration of monocytes to the atherosclerotic plaque [1]. Therefore, VCAM-1 expression can be correlated with the extent of plaque formation, making VCAM-1 an ideal biomarker for atherosclerosis [1, 3, 4]. VCAM-1 expression is also upregulated on the endothelium of rejected transplanted organs [3], and in breast cancer [5], lung cancer [6], and colorectal cancer [7]. Due to its role in a range of physiological and medical conditions, VCAM-1 has received considerable attention in the last decades.

VCAM-1 is a 90 kDa type I transmembrane glycoprotein and a member of the immunoglobulin (Ig) superfamily of proteins [8]. VCAM-1 interacts with cell surface proteins such as integrin α_1_β_4_ (VLA-4), one of the main binding partners, as well as other integrins and non-integrin proteins such as Galectin-3 (in eosinophils), and SPARC/Osteonectin, among others [1–3, 9–12]. VCAM-1 interacting proteins are critical during inflammation, as they promote leukocyte migration/transvasation [12–16]. VCAM-1 interaction with (some) protein ligands leads to downstream signaling resulting in cytoskeleton re-organization and loosening of cell-to-cell junctions, which facilitates leukocyte transvasation [17]. VCAM-1 is composed of 3-7 Ig-like domains, a single transmembrane domain, and a 19 amino acids cytoplasmic domain [1, 8, 18]. Human VCAM-1 is expressed as either 7 (7D) or 6 (6D) Ig-like domain containing forms, while mouse VCAM-1 has a 7D and a truncated 3D form [8, 18, 19]. In humans, the full-length 7D form, rather than the 6D form, is predominantly expressed [8, 20]. The 7D protein shows efficient binding to VLA-4 and is the primary mediator of cell adhesion [9, 21]. The 6D VCAM-1 does not contain domain 4, and shows less efficient cell adhesion, with decreasing efficiency of binding under increasing shear force [1, 21]. VCAM-1 can be proteolytically cleaved near the transmembrane domain by matrix metalloproteinases (MMPs), including the sheddase ADAM17 [22, 23], resulting in a soluble VCAM-1 version [24]. The level of soluble VCAM-1 6D form increases in certain pathologies such as RA and systemic lupus erythematosus [25, 26]). There is high sequence similarity between Ig-like domains 1 and 4, 2 and 5, and 3 and 6, due to a proposed intragene duplication event [18]. Of all duplicated domain pairs, domains 1 and 4 share the highest sequence similarity [27], and these domains are also involved in the interaction of VCAM-1 with VLA-4, expressed in leukocytes [8, 9, 28]. Domain 2 appears to be structurally required for domain 1 function, and specific residues in domain 2 are involved in the interaction with VLA-4 (α_1_β_4_) and, more importantly, with α_4_β_7_ [29, 30]. The interaction with Galectin-3, on the other hand, typically requires LacNAc disaccharides (Galβ1-4GlcNAc) found internally in *N*- or *O*-linked glycans [14], suggesting that human VCAM-1 could bind Galectin-3 through its Ig-like domains 3, 4, 5, and/or 6, which carry 1,1, 2, and 2 *N*-linked glycans, respectively. On the other hand, domain 6 of VCAM-1 appears to be a key target for TNFα-induced angiogenesis [31], and antibody blockade of domain 6 impairs leukocyte transmigration but not adhesion, as well as lung cancer cell migration [3, 6, 32]. To date, the functional roles of only some of VCAM-1 domains have been elucidated.

Due to the important role of VCAM-1 in a range of pathologies, several antibodies targeting VCAM-1 have been developed [1] (see **Supplementary Table S1**). Anti-VCAM-1 antibodies have been tested in mouse models for their effectiveness against asthma [33] and atherosclerosis [34, 35]. Further, there are promising *in vitro* studies that utilize radiolabeled anti-VCAM-1 antibodies to target and treat early-stage brain metastases [36]. In addition, diagnostic ELISA kits for VCAM-1 expression have been used in multiple clinical trials (clinicaltrials.gov). Clearly, VCAM-1 antibodies play important roles both in the laboratory and the clinic.

Identifying new antibodies against membrane targets can be challenging. Monoclonal antibodies (mAbs) are typically derived from single B cell/hybridoma clones [37, 38] or from phage display libraries [39]. One key aspect required for any successful antibody discovery campaign is antigen presentation. The antigen should ideally be presented to the polyclonal antibody pools as close to its native conformation as possible, which can be difficult in the case of membrane proteins due to their hydrophobic transmembrane domain(s) [40]. To ensure native conformation, membrane proteins can be expressed on the surface of cells or other membrane environments such as nanodiscs [41]. Phage display biopanning is an excellent *in vitro* platform for the discovery of new antibodies against both soluble and membrane-bound targets like VCAM-1, and several mAbs derived from phage display biopanning have been approved for clinical use [42, 43].

Discovering antibodies against VCAM-1 is important not only due to the potential medical applications of targeting VCAM-1, but also to understand the biological function of this important molecule. Since the late 1980s, several groups have published anti-VCAM-1 antibodies (see **Supplementary Table S1**). These antibodies bind VCAM-1 domains 1, 4, and 6, or within the 1-3 and 4-7 regions of the molecule (**Supplementary Table S1**), and have shed light on functional aspects of VCAM-1. However, specific functional contributions of some VCAM-1 Ig-like domains (such as domains 3, 5, and 7) and their association with cell binding and intracellular signal transduction remain understudied. To increase the toolbox of antibodies that recognize specific Ig-like domains of VCAM-1, we performed phage display biopanning using a naïve human phage display library [40] and panned against membrane-bound mouse 7D VCAM-1. We identified multiple scFv antibodies that bind Ig-like domains 2, 3, or 5. We show here that scFv antibodies binding Ig-like domains 2 and 3 significantly reduce *in vitro* binding of macrophages to activated endothelial cells. These results suggest that mVCAM-1 Ig-like domains 2 and 3 play important functional roles, that these antibodies could help obtain a clearer picture of the structure-function relationship between VCAM-1 Ig-like domains and VCAM-1 physiological roles, and that antibodies targeting these domains could have diagnostic and/or therapeutic potential.

## 2. Methodology

### 2.1. Plasmid Construction and Site-Directed Mutagenesis

All vectors used in this study are described in **Supplementary Table 2** and mutagenesis primers are described in **Supplementary Table 3**. The pUCIDT-Amp vector containing the *Mus musculus* VCAM-1 open reading frame (Uniprot accession ID P29533) codon optimized for expression in Chinese Hamster Ovary (CHO) cells was purchased from Integrated DNA Technologies (IDT) (See **Supplementary Material** for full mVCAM-1 sequence). mVCAM1 was subcloned from pUCIDT-Amp-VCAM-1 (LZCBI17) into pEGFP-N1 through NheI and BamHI double restriction digestion and ligation to generate full-length mVCAM1 C-terminally tagged with eGFP (LZCBI18).

mVCAM-1 Ig-like domain deletion mutants (LZCBI121-127 and LZCBI129-130) were constructed by deletion of 1 or more of the 7 Ig-like domains, similar to a previously published strategy [32]. The description of the mutagenesis strategy can be found in Supplementary Methods.

Phagemids from selected phage clones were extracted using Qiagen QIAprep Spin Miniprep kit (Qiagen, 27104) following the manufacturer’s protocol and sequenced as previously described [40, 41]. The DNA encoding the selected scFvs was subcloned from the phagemid vectors into pET28b(+)-LPETG vector (a modified Novagen vector that incorporates a C-terminal LPETG tag, as described in Supplementary information) through NcoI and NotI double restriction digestion and ligation. The scFv sequences obtained were reformatted into full length mouse IgG2a monoclonal antibodies (**Supplementary Table 2**) using the InFusion HD cloning kit (Takara, 639648), following previously published methodologies [41, 44].

All clones made in this study were sequenced to ensure the correct DNA sequence.

### 2.2. Mammalian Cell Culture Maintenance and Transfection

Chinese Hamster Ovary (CHO) XL99 and Human Endothelial Kidney (HEK) 293 cell lines were maintained in CD CHO medium (Gibco™, 10743029) or FreeStyle™ 293 Expression medium (Gibco™, 12338018), and supplemented with 0.4% (v/v) anti-clumping agent (Gibco™, 0010057DG) and GlutaMAX™ Supplement (Gibco™, 35050061). ExpiCHO cells were maintained on ExpiCHO medium as described by the manufacturer (Thermo Fisher, A29131). Cells were routinely grown at 37 °C, 7.5% CO_2_, and 125 rpm. Adherent CHO-K1 cells were maintained in Ham’s F-12K (Kaighn’s) Medium (Gibco™, 21127022) supplemented with 10% Fetal Bovine Serum (FBS) (Gibco™, 10099141) and incubated at 37 °C, 7.5% CO_2_.

CHO XL99 and HEK293 cells were transfected using a DNA:polyethyleneimine (PEI)-Max complex as previously described [41]. For transfection of CHO-K1 cells, 1.4 × 10^6^ CHO-K1 adherent cells were seeded into T25 flasks and incubated overnight at 37 °C, 7.5% CO_2_. The next day, pEGFP-N1-mVCAM-1 or pEGFP-N1 (negative control) plasmids were transfected into CHO-K1 cells using Lipofectamine™ 3000 Reagent (Invitrogen™, L3000001) and OptiMEM™ Reduced Serum Medium (Gibco™, 31985062) and following manufacturer’s procedure. ExpiCHO cells were transfected as per the manufacturer’s instructions using an ExpiFectamine CHO Transfection kit (Thermo Fisher, A29131). Expression of VCAM-1_eGFP and mutants was monitored using BioRad ZOE™ Fluorescent Cell Imager and by flow cytometry (see below).

Macrophage J774A.1 cells and SVEC4-10 endothelial cells (CRL-2181) were obtained from the American Type Culture Collection. The J774A.1 cells were maintained in a nontreated 100 × 20 mm cell culture dish (Corning®,430591) containing RPMI 1640 (Thermo Fisher, LTS11875093) with 10% fetal calf serum (Thermo Fisher, 10099141), penicillin (100 U/ml) (Thermo Fisher, 15140122), and 1% L-glutamine (Thermo Fisher, 25030081), and cultured in an incubator at 37 °C, 5% CO_2_. The SVEC4-10 cells were maintained in tissue culture treated T75 flask (Sarstedt®) containing DMEM medium (Gibco™, 11965092) supplemented with 10% fetal calf serum (Gibco™, 10099141), penicillin (100 U/ml), and 1% L-glutamine, and cultured in an incubator at 37 °C, 5% CO_2_.

### 2.3. Phage Display Biopanning

Cell-based phage display biopanning was performed following procedures previously described [40, 41, 45]. The scFv phage library used for the first round of biopanning was the Jones-Mahler human naïve library [40]. The rounds of biopanning alternated between CHO XL99 cells and HEK293 cells. GFP positive cells were sorted at the Queensland Brain Institute Flow Cytometry facility (UQ) using a BD FACSymphony^TM^ S6 Cell Sorter.

### 2.4. Flow Cytometry Analysis of Phage Pools

CHO XL99 cells were transfected with the pEGFP-N1-mVCAM-1 plasmid (**Supplementary Table S2**) and grown for 24 hours. 10^11^ phages from the naïve phage display library or round 1 and round 2 amplified phage pools were diluted in 1.8 mL of 2% skim milk in PBS (MPBS) and incubated at 4 °C, rotating for 20 minutes. A total of 10^6^ transfected and non-transfected CHO XL99 cells were harvested by centrifugation at 300 x *g* for 5 minutes and washed with 1 mL of PBS. Next, cells were incubated with each diluted phage pool in MPBS for 1 hour, rotating at 4 °C. Cells were washed 3 times with 1 mL PBS and were then incubated in a 1:200 dilution of mouse anti-M13 mAb (Sino Biological Cat# 11973-MM05T, RRID:AB_2857926) in MPBS on ice for 1 hour. Cells were harvested and washed twice with 1 mL PBS as above. Next, cells were incubated in a 1:400 dilution (1.25 μg/mL) of goat anti-mouse Dylight650 (Abcam Cat# ab96874, RRID:AB_10679531) in MPBS in the dark, on ice, for 1 hour. Cells were harvested and washed 3 times in PBS as above, and gently resuspended in 100 µL 4% paraformaldehyde (PFA) in PBS. After incubation on ice for 15-20 minutes, cells were washed 3 times as above and resuspended in 200 µL of PBS prior to analysis by flow cytometry using a Beckman Coulter CytoFLEX S (Beckman Coulter Life Sciences, Mount Waverley, VIC, Australia).

### 2.5. Whole-Cell Phage Binding ELISA

Individual phage clones from the second biopanning round were tested for binding to cell surface expressed mVCAM-1 by whole-cell Enzyme-Linked Immunosorbent Assay (ELISA). The phage clones were prepared in a 96-well format as previously described [40, 41], and were used for ELISA on 96-well plates coated with CHO-K1 cells expressing mVCAM-1-eGFP or eGFP alone (negative control). To coat the plates, transfected CHO-K1 cells were seeded onto 96-well plates to a density of 40,000 cells/well in 200 µL of F12K media supplemented with 10% FBS and Penicillin-Streptomycin antibiotic, and were incubated overnight at 37°C, 7.5% CO_2_. Next, cells were fixed in 4% PFA in PBS, and plates were left to dry at room temperature for 10 minutes. Phages and cells were separately blocked in MPBS for 1 hour at room temperature, and then the blocked phages were added to the blocked cells and incubated for 1 hour at room temperature. Plates were washed six times with PBS + 0.1% Tween 20 using a plate washer (BioTek, ELx405 Select). Next, a 1:5,000 dilution of mouse anti-M13 monoclonal antibody (Sino Biological Cat# 11973-MM05T, RRID:AB_2857926) in MPBS was added to the wells, followed by 1 hour room temperature incubation. Plates were washed as above, then incubated in a 1:5,000 dilution in MPBS of Goat Anti-Mouse IgG (H+L)-HRP conjugated secondary antibody (Bio-Rad Cat# 1706516, RRID:AB_2921252) for 1 hour at room temperature. After the final wash, plates were developed using TMB substrate (BioLegend, 423001) following the manufacturer’s instructions. Absorbance at 450 nm was measured using a microplate spectrometer (BioTek PowerWave XS2). The controls for this experiment included cells transfected with pEGFP-N1 empty vector and cells incubated only with primary and secondary antibodies but no phages. We considered an absorbance reading as positive if it complied with the following two rules: 1) the absorbance of the phage sample on VCAM-1-eGFP transfected cells was at least 4-times greater than the average signal of all the control wells in that same plate, and 2) the absorbance of the same phage sample on GFP only transfected cells was below 2-times the value of the average of the control wells in the corresponding plate (**Supplementary Figure S2**).

### 2.6. Flow Cytometry Analysis of Selected Phages and Purified Antibodies

Flow cytometry was used to verify the binding of selected phages, purified scFv-LPETG clones, and purified mAbs to mVCAM-1-eGFP. To test the phages, CHO XL99 cells were transfected or not with pEGFP-N1-mVCAM-1, and 24-48 hours later cells were aliquoted into wells of a round bottomed 96-well plate (2.5 x 10^5^ cells/well). Cells were pelleted by centrifugation at 300 x *g* for 5 minutes at room temperature, were washed with 200 µL of PBS, and were blocked with 25 µL of MPBS. The selected positive purified phages (30 µL) were blocked with 30 µL of 4% MPBS for 30 minutes at room temperature in a separate 96-well plate, and blocked phages were then combined with the blocked cells. After 1 hour incubation at 4 °C, cells were washed with 200 µL of PBS and incubated in 50 µL of 1:200 anti-M13 antibody (Sino Biological Cat# 11973-MM05T, RRID:AB_2857926) for 1 hour at 4 °C. Next, cells were washed 3-times with 200 µl PBS, and were incubated for 1 hour at 4 °C in 50 µl of 1:400 (1.25 μg/mL) goat anti-mouse IgG Dylight650 (Abcam Cat# ab96874, RRID:AB_10679531). Cells were washed as above, fixed in 4% PFA in PBS, and analyzed by flow cytometry as described above for the polyclonal flow cytometry.

A similar protocol was used for the scFv-LPETGs, except that 1 µg scFv was used per 10,000 cells instead of phages, the primary antibody used was a 1:200 dilution (2.5 μg/mL) of Purified Mouse anti-6XHIS with Control (BD Biosciences Cat# 552565, RRID:AB_394432), and no fixing with PFA was necessary.

For the mAbs, a 1:3 mixture of non-transfected to transiently transfected ExpiCHO cells were collected by centrifugation at 300 x *g* for 5 minutes and blocked in ice-cold FACSBlock (3% Fetal calf serum (FCS) (Gibco, A3160902), 0.1% bovine serum albumin (BSA) (Bovogen, SFBS-F), and 0.1% sodium azide) at a concentration of 1×10^6^ cells/ml for 30 minutes. Next, cells were harvested by centrifugation and resuspended in ice-cold FACSWash buffer (0.2% FCS, 0.1% BSA, 0.1% sodium azide) at a concentration of 5×10^6^ cells/mL, and 40,000 cells were aliquoted/well in a 96-V-shaped wells plate. The FACSWash buffer was removed by centrifugation at 400 x *g* for 5 minutes, and cells were resuspended in 50 µl of FACSWash containing 11.1 µg/mL of each reformatted anti-VCAM-1 antibody or isotype control mIgG2a (Miltenyi Biotec Cat# 130-106-546, RRID:AB_2661589), a 1:200 dilution (2.5 μg/mL) of rat anti-mouse VCAM-1 (Millipore Cat# CBL1300, RRID:AB_2214062), or 1:400 dilution of mouse IgG1 anti-FLAG (Cell Signaling Technology Cat# 8146, RRID:AB_10950495). For no antibody control wells, cells were incubated in FACSWash only. After 1 hour incubation at 4 °C, cells were collected by centrifugation and washed 3-times with 150-200 µl of FACSWash, followed by a 30-minute incubation at 4 °C in FACSWash containing 1:400 (1.25 μg/mL) goat IgG anti-mouse IgG H&L Dylight650 (Abcam Cat# ab96874, RRID:AB_10679531) or 1:200 (1 μg/mL) Goat anti-rat-PE-Cy7 (BioLegend Cat# 405413, RRID:AB_10661733) antibodies, or FACSWash for no antibody controls. Next, cells were collected by centrifugation, washed 3-times in FACSWash, and resuspended in 100 µl of FACSWash supplemented with 0.25% PFA. Cells were analyzed by flow cytometry using a Beckman Coulter CytoFLEX S and the data collected was analyzed using CytExpert and FlowJo v10.6.2 (FlowJo LLC, BD Biosciences). Median fluorescence intensity (MFI) was used as a measure of binding of the different mAbs to VCAM-1 and its Ig-like mutant versions.

### 2.7. Expression and Purification of scFv and Full-Length Antibodies

scFv-LPETG antibodies were expressed in SHuffle^®^ Express *E. coli* cells (New England BioLabs® Inc.). Overnight cultures of the SHuffle strains carrying the different pET28b-scFvs were diluted 1:100 in 200 mL TB medium (Merck, T0918) supplemented with Kanamycin (30 µg/mL) and were grown at 37 °C at 200 rpm until cells reached log phase (OD_600_ = 0.8-1.0). The culture was allowed to reach room temperature, and scFv-LPETG expression was induced by adding 1 mM of Isopropyl β-D-1-thiogalactopyranoside (IPTG). Protein expression was conducted overnight at 25 °C, 200 rpm. Cells were harvested the following day by centrifugation at 5,000 x *g* for 10 minutes at room temperature. Bacterial pellets were resuspended in 30 mL of ice-cold equilibration buffer (20 mM sodium phosphate, 0.5 M NaCl, 20 mM imidazole, pH 7.4) by vigorous pipetting, and were sonicated on ice (SONICS Ultrasonic Processor VC-505) for 5 minutes (20% Amplitude, Pulse 1s on 1s off, 3 mm tip diameter). The lysate was incubated at 4 °C, rotating, for 1 hour and then centrifuged (Avanti JXN-30) at 18,000 x *g* for 30 minutes at 4 °C. The supernatant was recentrifuged at 20,000 x *g* for 30 minutes at 4 °C, transferred to a pre-cooled vessel, and the scFv-LPETGs were purified using the Äkta Explorer FPLC (Fast Protein Liquid Chromatography) system using a 1 mL HisTrap column (HisTrap™ High Performance, Cytiva, 17524701), and following standard procedures. The scFv-LPETG clones were eluted in elution buffer (20 mM sodium phosphate, 0.5 M NaCl, 400 mM imidazole, pH 7.4) on ice and dialyzed in 14 kDa cutoff dialysis tubing against a solution of 1X PBS overnight at 4 °C. The pH of the dialysis buffer was adjusted to 1 unit above the theoretical pI of each scFv (**Supplementary Table 4**). The dialyzed samples were concentrated using a centrifugal concentrator (Vivaspin® 6 Centrifugal Concentrator, VS0602) with a 10 kDa Molecular weight cut-off (MWCO) by centrifuging at 4,000 x *g* at 4 °C. The concentration of scFv-LPETG was determined with a Nanodrop Spectrophotometer, considering the calculated extinction coefficients (**Supplementary Table 4**). Samples were transferred to 1.5 mL Protein LoBind® tubes (Eppendorf, 0030108442) and stored at 4 °C for the short term or supplemented with 10% v/v final concentration of glycerol, frozen in liquid nitrogen, and stored long term at -80 °C.

mAbs were expressed as previously described [41] using ExpiCHO cells and the high titer protocol from the ExpiFectamine CHO Transfection kit (Thermo Fisher, A29131). The supernatant was loaded into a prepacked HiTrap MabSelect column (Cytiva, 28408253) and purified using an Äkta Explorer FPLC system. Purified mAbs were buffer exchanged and concentrated using Amicon Ultra Centrifugal filters and stored in 10% glycerol, PBS pH 7.4 at -80 °C until use.

### 2.8. SDS-PAGE and Western Blotting

The presence of antibodies in the supernatant and eluates was determined using sodium dodecyl sulphate–polyacrylamide gel electrophoresis (SDS-PAGE) as follows. For scFv-LPETGs, 5 µg of each purified antibody samples were mixed with 4X NuPAGE loading buffer supplemented with 10 mM DTT, heated at 95 °C for 5 minutes, centrifuged for 1 minute at 21,000 x *g*, and loaded in 10% acrylamide gels (Bio-Rad, 4561033). Protein detection was achieved by either staining the gel with SimplyBlue™ SafeStain (LC6065) following the manufacturer’s protocol, or by western blot. For purified mAbs, 5 µg of each purified antibody was resuspended on 4X SDS-PAGE loading buffer (200 mM Tris-HCl pH 6.8, 8% SDS, 0.4% bromophenol blue, 40% glycerol) supplemented or not with a final concentration of 10 mM DTT. The sample was heated at 95 °C for 5 minutes, centrifuged for 1 minute at 21,000 x *g*, and separated in a 4-15% stain-free gel (BioRad, 4568081). All gels were visualized on a BioRad Chemidoc imaging system.

For western blotting, proteins were transferred from acrylamide gels into a Polyvinylidene fluoride (PVDF) membrane using the BioRad Trans-Blot Turbo RTA Transfer Kit (1704272) and a Trans-Blot Turbo Transfer system (7 minutes at 1.3A), following the manufacturer’s protocol. The membrane was blocked with 5% skim milk in Tris buffered saline (50 mM Tris-Cl, pH 7.5, 150 mM NaCl) with 0.1% Tween 20 (TBS-T) for 1 hour at room temperature, rocking. The membrane was then incubated with a 1:5,000 dilution of anti-His-HRP (Miltenyi Biotec Cat# 130-092-785, RRID:AB_1103231) in 2% skim milk TBS-T overnight, rocking at 4 °C. After 3 washes with TBS-T for 5 minutes at room temperature, membranes were developed with Super signal West Dura Extended Duration Substrate kit (ThermoFisher Scientific, 34075) and visualized in the BioRad ChemiDoc gel imaging system.

### 2.9. Bioinformatic Analysis of scFv Sequences

To identify variable domains of immunoglobulins in the scFv sequences, we used IgBlast (IgBlast tool (nih.gov)) from the National Centre for Biotechnology Information (NCBI). To annotate the variable regions with the ImMunoGeneTics (IMGT) numbering system, we used Antigen receptor Numbering and Receptor Classification (ANARCI) by SAb-Pred (http://opig.stats.ox.ac.uk/webapps/newsabdab/sabpred/anarci/). The molecular weight, extinction coefficient, and theoretical pI for each scFv-LPETG (**Supplementary Table S4**) were calculated using ProtParam (<u>ExPASy - ProtParam tool</u>) based on the scFv’s amino acid sequences. Amino acid sequences were aligned using Clustal Omega (https://www.ebi.ac.uk/Tools/msa/clustalo/), which was also used to generate phylogenetic trees.

### 2.10. In Vitro Cell Adhesion Assay

SVEC4-10 endothelial cells were seeded into a 96-well plate at a density of 10,000 cells/well and incubated for 24 hours. The cells were then separately treated with 100 ng/mL lipopolysaccharide (LPS) (Sigma-Aldrich, L2630) for 24 hours. J774A.1 macrophages were stained on the same day with 25 ng/mL of DiOC6 for 24 hours at 37 °C. After 24 hours LPS stimulation, SVEC4-10 endothelial cells were treated with the following different antibodies at a concentration of 20 µg/mL, for 1 hour at 37 °C: rat anti-VCAM-1 (Millipore Cat# CBL1300, RRID:AB_2214062) was used as positive control; a random scFv (not mVCAM-1 binder, a kind gift from the National Biologics Facility, UQ) was used as negative control, and 6 different purified scFv antibodies from this work (1A9, 2D3, 2D8, 2E2, 2E6, and 3H4). After incubation, endothelial cells were washed once with PBS. The DiOC6-labelled J774A.1 cells were detached by washing with PBS and centrifuged at 500 x *g* for 5 minutes, followed by resuspension in DMEM, high glucose medium (Cat #11965-092, Thermo Fisher) supplemented with 10% Fetal Bovine Serum (Cat #26140079, Thermo Fisher) and 1% Penicillin-Streptomycin (Cat #15140122, Thermo Fisher). Next, DiOC6-labelled J774A.1 cells were added to SVEC4-10 endothelial cells at a density of 10,000 cells/well. After 10 minutes of coculture treatment, all wells were washed once with PBS and then fixed with 4% PFA in PBS for analysis. Fluorescent images were taken with an OLYMPUS CKX53 fluorescent microscope with the Cool LED pE-300 light source and OLYMPUS-DP74 camera. The relative fluorescent intensity was quantified with ImageJ. The experiment was performed twice with up to five replicates/group, for a total of 9.

### 2.11. Statistical Analysis of Data

Data are presented as Mean +/- standard error unless specified in the figure legend. One-way ANOVA followed by post-hoc Tukey test for pairwise comparisons was used in the analysis of significant differences in the cell adhesion assay. A *p* value < 0.05 was considered significant. Graphs were plotted using GraphPad Prism 10.

## 3. Results

### 3.1. Identification of Binders to mVCAM-1 Through Phage Display Biopanning

To identify scFv binders to mouse 7D VCAM-1 (mVCAM-1), we used a human naïve phage display library [46] and we panned against full length mVCAM-1 transiently expressed in mammalian cell lines. Whole-cell phage display biopanning allows expression of VCAM-1, a transmembrane glycoprotein with multiple disulphide bonds, in its native conformation with adequate post-translational modifications [40]. The full open reading frame (ORF) of the 7 Ig-like domain mVCAM-1 was fused with a C-terminal eGFP protein to monitor transfection efficiency, observe mVCAM-1 subcellular localization, and for effective cell sorting through FACS. Importantly, the VCAM-1 C-terminus is intracellular, and we expected no impact of the independently folding eGFP protein on VCAM-1 biosynthesis and trafficking. Indeed, VCAM-1-eGFP expressed and trafficked to its expected plasma membrane localization with efficiency (**Figure 1D-F** and **Supplementary Figure S1**). We performed whole cell biopanning using the Jones-Mahler human naïve scFv library (JM library) [46], and alternating mammalian cell lines (CHO XL99 cells for round 1 and HEK293 suspension cells for round 2). Alternating cell lines during cell-based biopanning facilitates depletion of irrelevant cell surface binders due to the differences in surface proteome among the different cell lines, thereby improving enrichment of relevant binders [40]. To check for enrichment of phages against mVCAM-1 in our phage pools and select a phage pool for screening, we performed polyclonal flow cytometry on the original library and the sub libraries **(Figure 1)**. We looked for an increase in the % of GFP+/Phage+ events (top right quadrant of the graphs, **Figure 1**) in transfected cells, indicating phage binding to cells with high mVCAM-1-eGFP expression **(Figure 1D-F)**. The JM library did not show specific phage binding to mVCAM-1 **(Figure 1D**, 1.75% of events in GFP+/Phage+ quadrant) or non-transfected cells **(Figure 1A**, 0.04% of events in Phage+ quadrant), as expected. Round 1 and 2 phage pools showed increased unspecific cell surface binding to untransfected cells, also as expected since phage binding cell surface mammalian proteins will also be enriched in this method **(Figure 1B-C**, 15.3% and 22% events in GFP-/Phage+ quadrant, respectively). However, we observed a clear increase in mVCAM-1 phage binding in Round 2 phage pool compared to JM library and Round 1 phage pools **(Figure 1D-F**, 1.75% *vs* 3.29% and 27.9% events in GFP+/Phage+ quadrant, respectively), with some background binding observed, as expected (**Figure 1D-F**, 0.9%, 5.73%, and 10.1% events in GFP-/Phage+ quadrant). This result indicated that the biopanning campaign was successful, and that we had effectively enriched for anti-VCAM-1 binding phages after two rounds of panning. To note, generally, 3 or 4 rounds of biopanning are required for enrichment of scFv-phages against a target; however, we obtained a clear enrichment of the library on Round 2, a result likely associated with the quality and size of the antigen and its presentation format (**Figure 1D-F**). Performing further rounds of biopanning could have led to the isolation of phages with higher affinity/avidity for mVCAM-1, but this also risked reducing sequence variability in isolated phages and our chances of isolating binders to different regions of mVCAM-1 (a key goal of this work).

**Figure 1.**
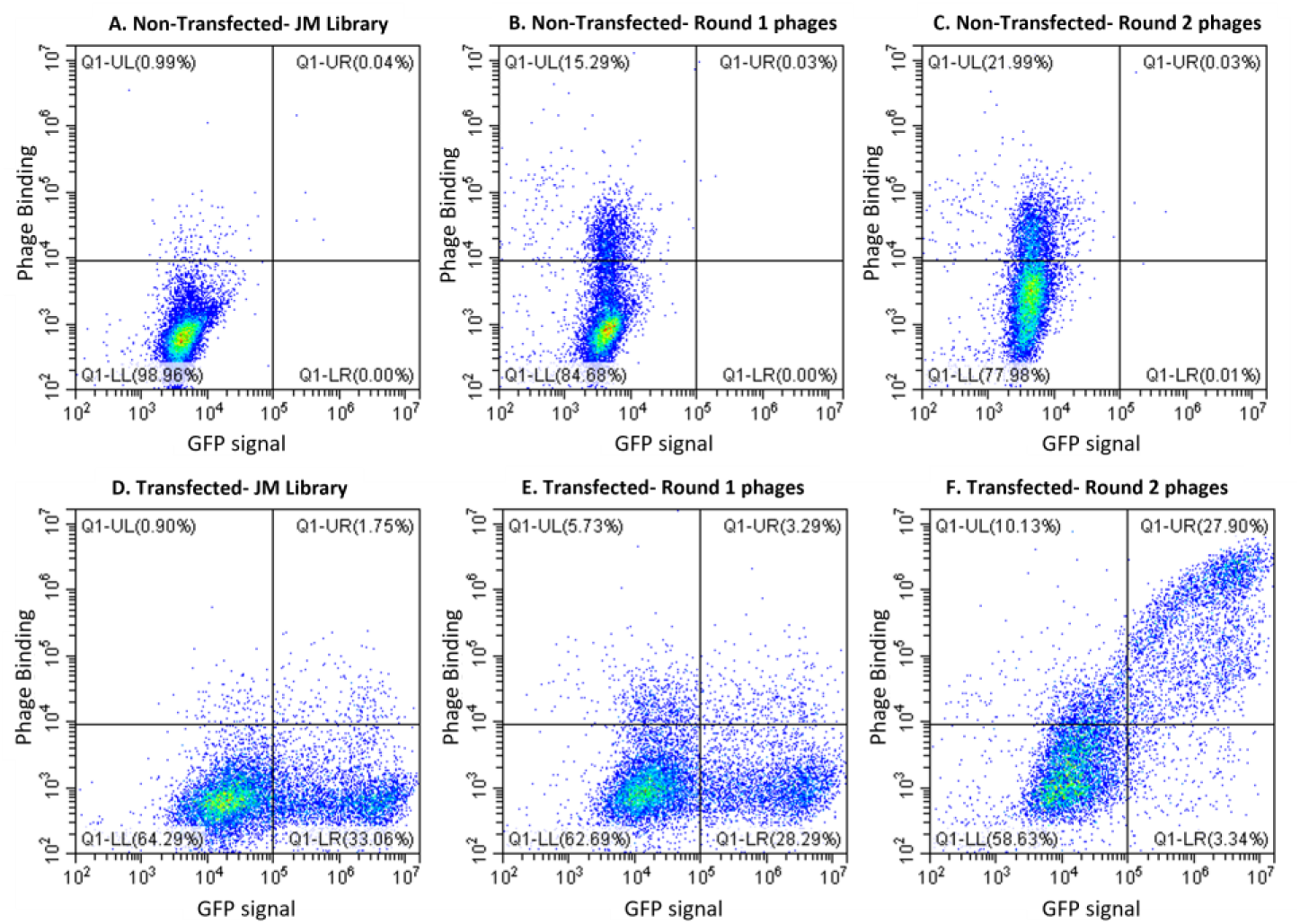
Flow cytometry analysis of the phage pools showed enrichment of anti-VCAM-1 phages in the round 2 phage pool. CHO XL99 cells non-transfected (A-C) or transfected with pEGFP-N1-mVCAM-1 (D-F) were incubated with the Jones-Mahler human naïve phage library (JM library) (A,D), Round 1 phage pool (B,E), or Round 2 phage pool (C,F). The plots depict GFP expression (x-axis) *vs* phage binding (y-axis).

To identify specific mVCAM-1 phage binders we first screened a total of 262 phage clones from the round 2 phage pool by monoclonal whole cell ELISA and identified several positive clones (∼ 6% positive clones) **(Supplementary Figure S2**). Next, we sequenced the scFv encoding DNA of the positive clones to identify unique clones. Multiple sequence alignment of the translated amino acid sequences of the clones identified 8 unique scFv sequences (**Supplementary Figure S2**, identified with numbers). The sequences for clones 1A9 and 2E6 were identified multiple times, while the other 6 sequences were represented only once. The coding regions of the scFvs were confirmed by IgBlast (IgBlast tool (nih.gov). Clones 1A9 and 2B9 had a kappa light chain, while the other 6 clones had a lambda light chain. Clones 2D3 and 3H4 shared the highest total sequence similarity among all 8 clones (**Supplementary Figure S3, left**), with identical complementary determining regions (CDR) 1 and 2 in the heavy chain as well as high total sequence similarity in their light chains (**Supplementary Figure S3, right**). Clones 2E2 and 2E6 shared an almost identical lambda light chain (**Supplementary Figure S3, right**). Clones 2D8 and 1A9 had identical CDR2 in the heavy chain, but all other CDR sequences differ (data not shown). All 8 clones had a unique CDR3 in the heavy chain (**Supplementary Figure S3, middle**), which is one of the most critical determinants of antibody specificity [47]. Together, these results suggest that each individual scFv likely binds to different mVCAM-1 epitopes and/or with varying affinities.

We used flow cytometry to confirm that all positive phage clones bound to mVCAM-1-eGFP transiently expressed on the surface of CHO XL99 cells (**Figure 2**, red). Some background binding to non-transfected cells was observed for some clones (2D8, 2E2, and 3H4), possibly due to the “sticky” nature of the phages and slight differences in sample processing (**Figure 2**, blue). We observed two different binding patterns for the phages (**Figure 2**): while most phages followed a linear relationship between the phage signal and GFP signal that eventually saturated and plateaued, some phages such as 2E2 and 2D3 showed a biphasic curve in which higher mVCAM-1-eGFP expression was required to observe phage binding (**Figure 2D,F)**. This result may suggest differences in binding strength for mVCAM-1 by the different phage clones (possibly due to differences in affinity of the scFvs for VCAM-1 or differences in the number of scFvs displayed on the phage surface), or differences in epitope availability. In summary, we performed two rounds of whole cell biopanning using a human naïve phage display library and mammalian cells transiently expressing mVCAM-1, and isolated 8 unique scFv sequences that bind the 7D mVCAM-1 isoform.

**Figure 2.**
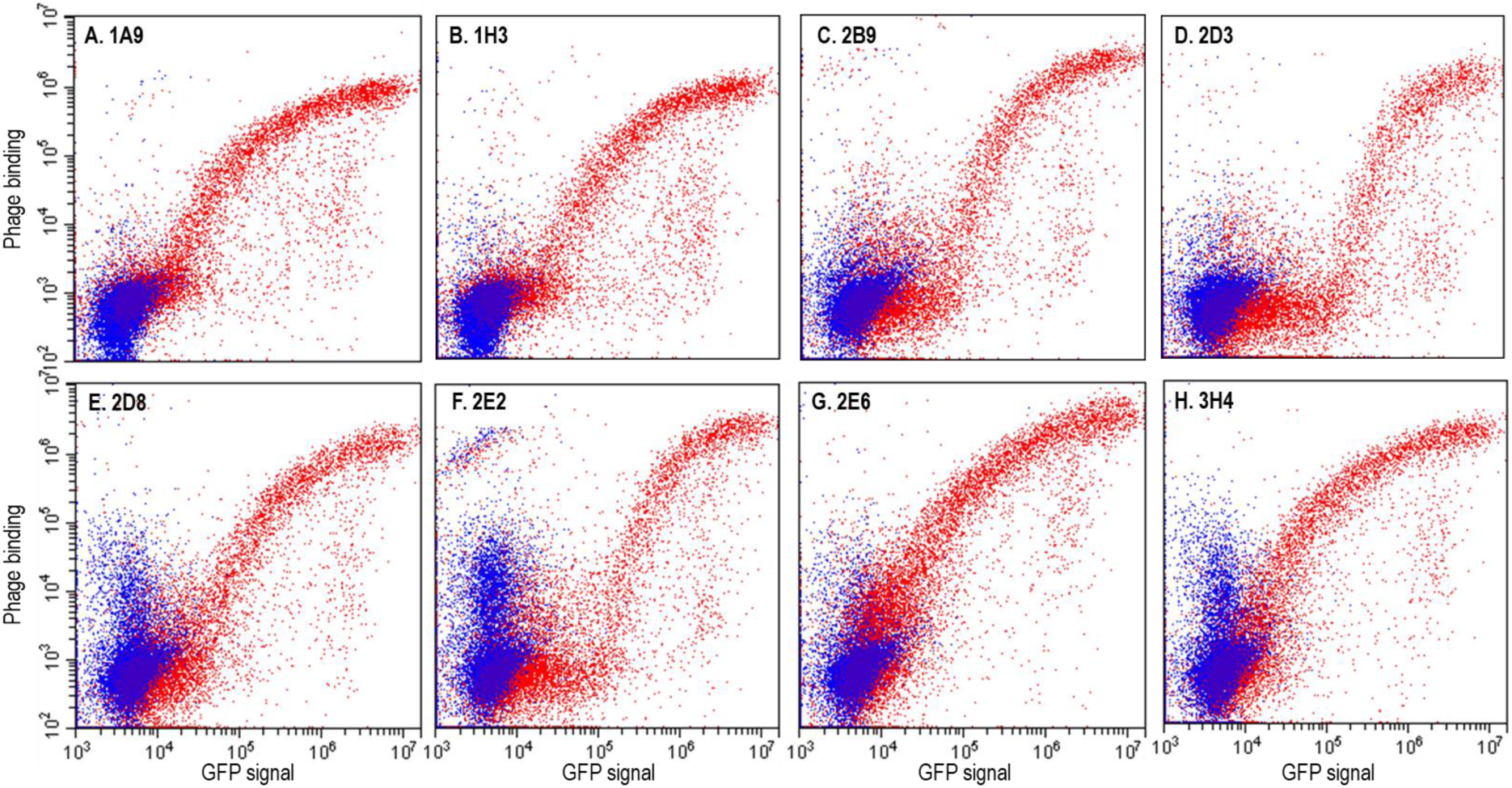
Monoclonal flow cytometry confirmed the positive phage clones bound to mVCAM-1. Positive phage clones from round 2 phage pool selected by whole-cell ELISA were tested for binding to mVCAM-1 by flow cytometry using CHO XL99 cells transfected with mVCAM-1-eGFP (red) or not (blue). Shown are the 8 unique clones selected (**A**) 1A9, (**B**) 1H3, (**C**) 2B9, (**D**) 2D3, (**E**) 2D8, (**F**) 2E2, (**G**) 2E6, and (**H**) and 3H4. The plots depict GFP expression (x-axis) *vs* phage binding (y-axis).

During DNA sequence analysis we also noticed that the amino acid sequence of clones 1H3 and 2B9 contained undesired sequence liabilities. Clone 1H3 had two Cys residues located within its CDR3-H which could lead to alterations in antibody binding specificity due to non-native disulfide bond formation and/or folding defects. On the other hand, clone 2B9 had an *N*-linked glycosylation sequon in CDR2-H which could be occupied following expression in eukaryotic cells. The presence of an *N*-linked glycan could impact folding, expression, and/or binding of 2B9 antibodies to the target. Therefore, the 1H3 and 2B9 clones were not further pursued.

### 3.2. Verifying Expression and mVCAM-1 Binding for all Unique scFv-LPETGs

The other six unique clones were subcloned into a pET28b(+)-LPETG vector. The LPETG tag is a sequence of 5 amino acids that can be used for sortase-mediated antibody conjugation [48, 49], allowing the construction of, for example, antibody drug conjugates and other chimeras. The tag is located exactly between the end of the scFv sequence and the LEHHHHHH C-terminal 6xHis tag, with no additional sequences, and its presence did not impact expression compared to scFvs alone (data not shown). These scFv-LPETG antibodies were expressed in *E. coli* SHuffle, a strain that promotes the formation of disulphide bonds in the cytoplasm, which are essential for scFv antibodies. The size of the scFv-LPETGs was calculated by ProtParam to be ∼ 27 kDa (**Supplementary Table S4**), as expected for scFvs. Expression of the scFv-LPETGs in *E. coli* SHuffle cells was followed by protein extraction and purification using a HisTrap column, dialysis, and concentration (**Supplementary Figure S4**). All scFv-LPETGs were successfully expressed and purified from bacteria.

To verify that the purified scFv-LPETG antibodies bound mVCAM-1 when not associated with a phage particle, we again performed flow cytometry (**Figure 3**). Non-transfected cells (negative control), and cells transfected with mVCAM-1-eGFP were incubated with purified scFv-LPETGs, and then stained with mouse anti-His antibody followed by anti-mouse IgG-Dylight650 antibody. As expected, all unique scFv-LPETGs showed binding to mVCAM-1, with only minor background binding of some of the antibodies to untransfected cells (**Figure 3**). The linear *vs* biphasic cell binding patterns previously observed for the phages were still observed with the purified scFv-LPTEG antibodies, indicating that this is a property of the scFv and not of the phage-bound scFv clones (**Figure 2D,F**, **Figure 3B,D**, clones 2D3 and 2E2). In conclusion, all selected scFvs were expressed in bacteria and bound mVCAM-1 expressed in CHO cells in the native membrane environment.

**Figure 3.**
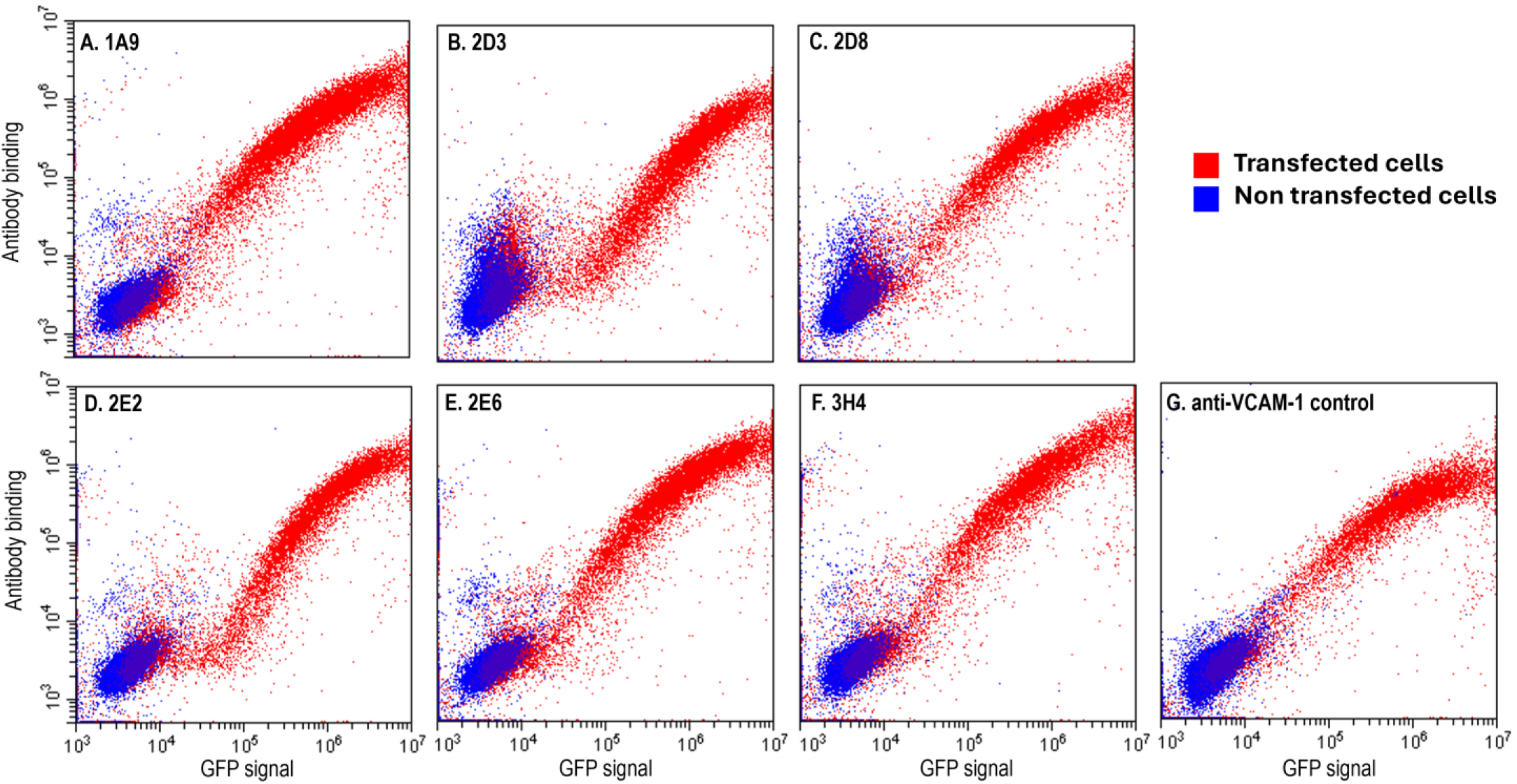
Purified scFv-LPETGs bind mVCAM-1. ExpiCHO cells transiently transfected with mVCAM-1-eGFP (red) or not (blue) were stained with purified scFv-LPETGs. The scFvs were detected with a mouse anti-HIS antibody and goat anti-mouse Dylight650. Shown are the 6 unique clones (**A**) 1A9, (**B**) 2D3, (**C**) 2D8, (**D**) 2E2, (**E**) 2E6, and (**F**) 3H4, and (**G**) anti-VCAM-1 CBL1300 as positive control. The plots depict GFP expression (x-axis) *vs* antibody binding (y-axis).

### 3.3. Ig-Like Domain Antibody Binding Mapping

VCAM-1 has 7 Ig-like domains, some of which have been associated with different functions (**Supplementary Table S1**). To determine to which mVCAM-1 Ig-like domain the discovered antibodies bind, we generated a suite of mVCAM-1 mutants in which up to 4 different Ig-like domains were simultaneously deleted from mVCAM-1 (as depicted in **Figure 4A**). Although all constructs were eGFP tagged, it was possible that some of the mutations led to lower cell surface expression while not substantially reducing intracellular GFP signal (**Supplementary Figure S5**). To ensure that all the deletion mutants were expressed on the cell surface, we inserted an N-terminal FLAG tag between the signal sequence and the first Ig-like domain in all constructs and performed flow cytometry using an anti-FLAG antibody (**Figures 4A,B**). Addition of the FLAG-tag did not appear to negatively influence the cell surface expression levels of the wild-type (WT) construct (compare anti-VCAM-1 binding to the untagged and FLAG-tagged WT constructs, **Figure 4A,B, Supplementary Figure S6**). As expected, the anti-FLAG antibody bound to all constructs (**Figure 4B, Supplementary Figure S6**), demonstrating that all constructs were expressed on the cell surface. However, the Δ5-7, Δ6-7, and Δ7 constructs showed weaker anti-FLAG binding when compared to the WT construct, with the Δ5-7 construct being the most impacted (**Figure 4B, Supplementary Figure S6**), indicating lower cell surface expression of these mutants. This result was not due to lower overall construct expression, since GFP signal in the cells transfected with the mutants was similar or higher than in the WT controls (**Supplementary Figure S5**). Further, since the Δ5 construct showed similar or higher anti-FLAG binding than the Δ6-7 and Δ7 constructs (**Figure 4B, Supplementary Figure S6**), it is not the deletion of domain 5 that led to the substantially lower expression levels in the Δ5-7 mutant (**Figure 4B**). While the lower expression of the Δ5-7 mutant could impact the ability of the antibodies to recognize the remaining Ig-like domains on this construct, the availability of the other well-expressed constructs was enough to provide certainty regarding the location of the (main) antibody binding site in the mVCAM-1 molecule.

**Figure 4.**
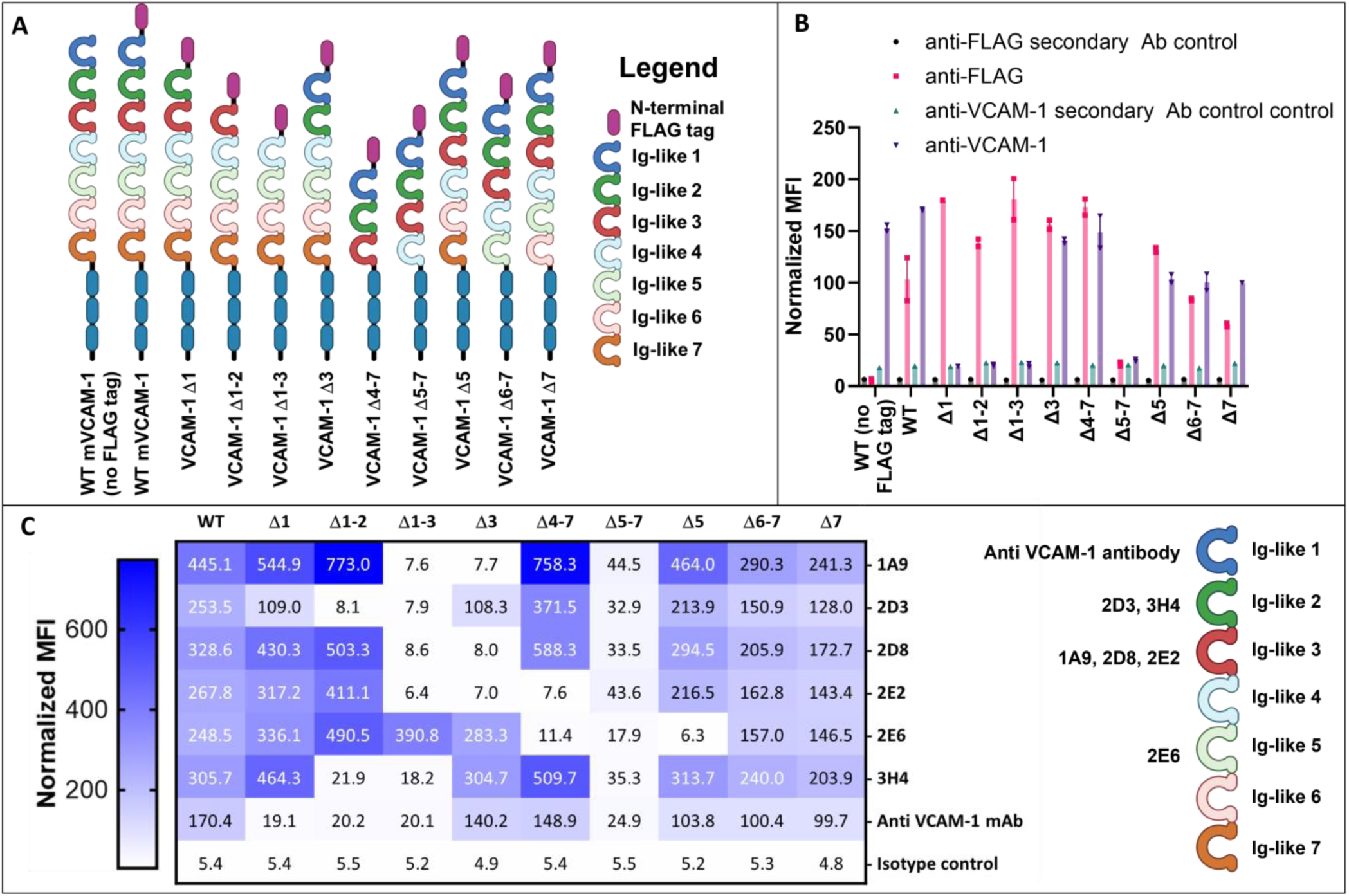
Ig-like domain mapping of the isolated antibodies. **(A)** Schematics of mVCAM-1 Ig-like domain deletion mutants used in this study. **(B, C)** Flow cytometry analysis of the binding of the commercial anti-mVCAM-1 antibody (CBL1300) **(B, C)**, anti-FLAG **(B)**, and our new full-length IgG2a reformatted antibodies **(C)** to cells transiently transfected with mVCAM-1-eGFP WT (with or without N-terminal FLAG) or FLAG-tagged Ig-like deletion constructs. Overall VCAM-1 cell surface expression was determined using anti-FLAG antibody or anti-VCAM-1 CBL1300 antibody **(B)**. CBL1300 required a different secondary antibody control than the FLAG antibody, therefore results with its own secondary antibody-only control is also shown (**B**). **(C)** Table showing the normalized MFI values for each of the VCAM-1/antibody combinations (also see **Supplementary Figure S8**). The color scheme correlates with MFI value, with lighter colors indicating lower antibody binding and darker colors higher antibody binding. On the right, a schematic representation of the mVCAM1 Ig-like domain structure showing the (tentative) binding location for the different antibodies tested in this study. Displayed is the average normalized MFI-Max-Min (mVCAM-1-eGFP-positive, antibody positive cells normalized to MFI of the mVCAM-1-eGFP-negative, antibody negative cells) from one experiment with 2 replicates. Schematics were created with BioRender.com.

Flow cytometry analysis of the CBL1300 (M/K2) rat anti-VCAM-1 antibody showed strong binding to the WT and the Δ4-7, Δ6-7, and Δ7 constructs (**Figure 4C**, **Table 1, Supplementary Figure S6**). Binding of this antibody to the Δ5-7 construct was poor, similar to what was observed with the anti-FLAG antibody (**Figure 4B-C, Supplementary Figure S6**). The commercial antibody did not bind the Δ1, Δ1-2, and Δ1-3 constructs, showing similar binding as the secondary antibody only control for those mutants (**Figure 4B, Supplementary Figure S6**). These results indicate that the anti-VCAM-1 commercial antibody binds to Ig-like domain 1 in mVCAM-1, as previously shown (**Supplementary Table S1**), validating our experimental design.

For this flow cytometry experiment we used the full-length mouse IgG2a reformatted versions of the scFvs (**Supplementary Figure S7**). As we hoped, the results demonstrate that the antibodies bound to different Ig-like domains (**Figure 4C, Supplementary Figure S8**). Antibodies 1A9, 2D8, and 2E2 appeared to bind the Ig-like domain 3 of mVCAM-1 (**Figure 4C, Supplementary Figure S8**), since they were able to bind to all constructs except for the Δ1-3, Δ2-3 and Δ3 constructs (**Figure 4C, Supplementary Figure S8**). Interestingly, antibody 2E2 also showed reduced binding to the Δ4-7 construct, suggesting that the binding site may include Ig-like domain 4, or may be affected by its deletion due to a conformational/structural change or to altered availability due to enhanced proximity to the cell membrane of the Δ4-7 construct (**Figure 4C, Supplementary Figure S8**). On the other hand, antibodies 2D3 and 3H4 appeared to bind the Ig-like domain 2, since they were able to bind all constructs except for the Δ1-2 and Δ1-3 constructs (**Figure 4C, Supplementary Figure S8**). To note, however, 2D3 also showed somewhat lower binding relative to the WT construct for the Δ1 mutant, suggesting that the binding site for 2D3 may be close to and/or involve a portion of the Ig-like domain 1 or connecting region, or alternatively that the absence of domain 1 impacts the binding site of 2D3 (**Figure 4C, Supplementary Figure S8**). Interestingly, as discussed above, sequence alignment of 3H4 and 2D3 showed that they share an almost identical heavy chain variable region (except for CDR3-H), and the highest similarity in light chains among all analyzed scFvs (**Supplementary Figure S3**), which could explain the similar, but not identical binding to mVCAM-1. Finally, the 2E6 antibody appeared to bind the Ig-like domain 5, since it was able to bind all constructs except for the Δ4-7, Δ5-7, and Δ5 constructs (**Figure 4C, Supplementary Figure S8**). Similar to the results with the anti-FLAG and commercial anti-VCAM-1 antibodies, none of our antibodies was able to bind well to the Δ5-7 mutant (**Figure 4C, Supplementary Figure S8**), probably due to the poor cell surface expression and potential associated folding issues in this construct (see above). We note that it is possible that some or all of the antibodies tested here are also able to bind (albeit less strongly) to the corresponding duplicated, yet not identical, Ig-like domain (i.e. 2 and 5, and 3 and 6 [18]). However, potential binding to the duplicated Ig-like domain was not observed in this experiment, either because of low sequence similarity at the specific epitopes or due to the signal being below the limit of detection for this assay. In conclusion, we were able to isolate multiple antibodies that bind to different Ig-like domains of mVCAM-1, including Ig-like domains 2, 3, and 5 (**Figure 4C, Supplementary Figure S8**).

### 3.4. scFv Antibodies Against VCAM-1 Ig-Like Domains 2 or 3 Can Block Macrophage Attachment to Endothelial Cells

Blocking different Ig-like domains in VCAM-1 with antibodies can lead to different functional impacts, depending on where the antibody binds, and this has been clearly shown for Ig-like domains 1, 4, and 6 [1, 8, 9, 28, 32] (**Supplementary Table S1**). One of the most important physiological roles of VCAM-1 is to facilitate macrophage recruitment into an inflammatory site [1]. Here, we tested if the antibodies we identified had an impact on the binding of macrophages to activated endothelial cells natively expressing 7D mVCAM-1. We performed this analysis using the scFv-LPETGs instead of the full-length antibodies to reduce the possibility of steric hindrance of the full-length antibody on mVCAM-1/ligand interactions, and thereby increase certainty of a direct impact of Ig-like domain blocking on the attachment phenotype. To do this, SVEC4-10 endothelial cells were treated with LPS for 24 hours to induce mVCAM-1 cell surface expression [50]. Next, different scFv-LPETGs were supplemented to the medium and cells were incubated for 2 hours to block mVCAM-1. Then, DiOC6-labelled macrophages were cocultured with endothelial cells for 10 minutes. Green fluorescence on the cell culture layer after washes represent macrophages that remained attached to the endothelial cells. As expected for a full-length antibody binding VCAM-1 Ig-like 1 domain (**Figure 4, Supplementary Table S1**), treatment with the commercial anti-mVCAM-1 significantly reduced macrophage ability to attach to endothelial cells compared to the non-antibody control, as demonstrated by a significant reduction in green fluorescence intensity (**Figure 5**, p < 10^-4^). Also as expected, the non-anti-VCAM-1 scFv antibody control did not significantly alter macrophage attachment, and its MFI value was similar to the non-antibody control (**Figure 5**). Interestingly, all anti-VCAM-1 scFv-LPETGs reduced macrophage attachment to endothelial cells and, with the exception of 2E6 (p = 0.056), the other 5 scFv-LPETG antibodies significantly reduced macrophage attachment when compared to the non-antibody treated control (**Figure 5**). Together, the results show that scFv antibodies able to bind domains other than 1, 4, and 6 [1, 8, 9, 28, 32] can also reduce macrophage attachment to endothelial cells *in vitro*.

**Figure 5.**
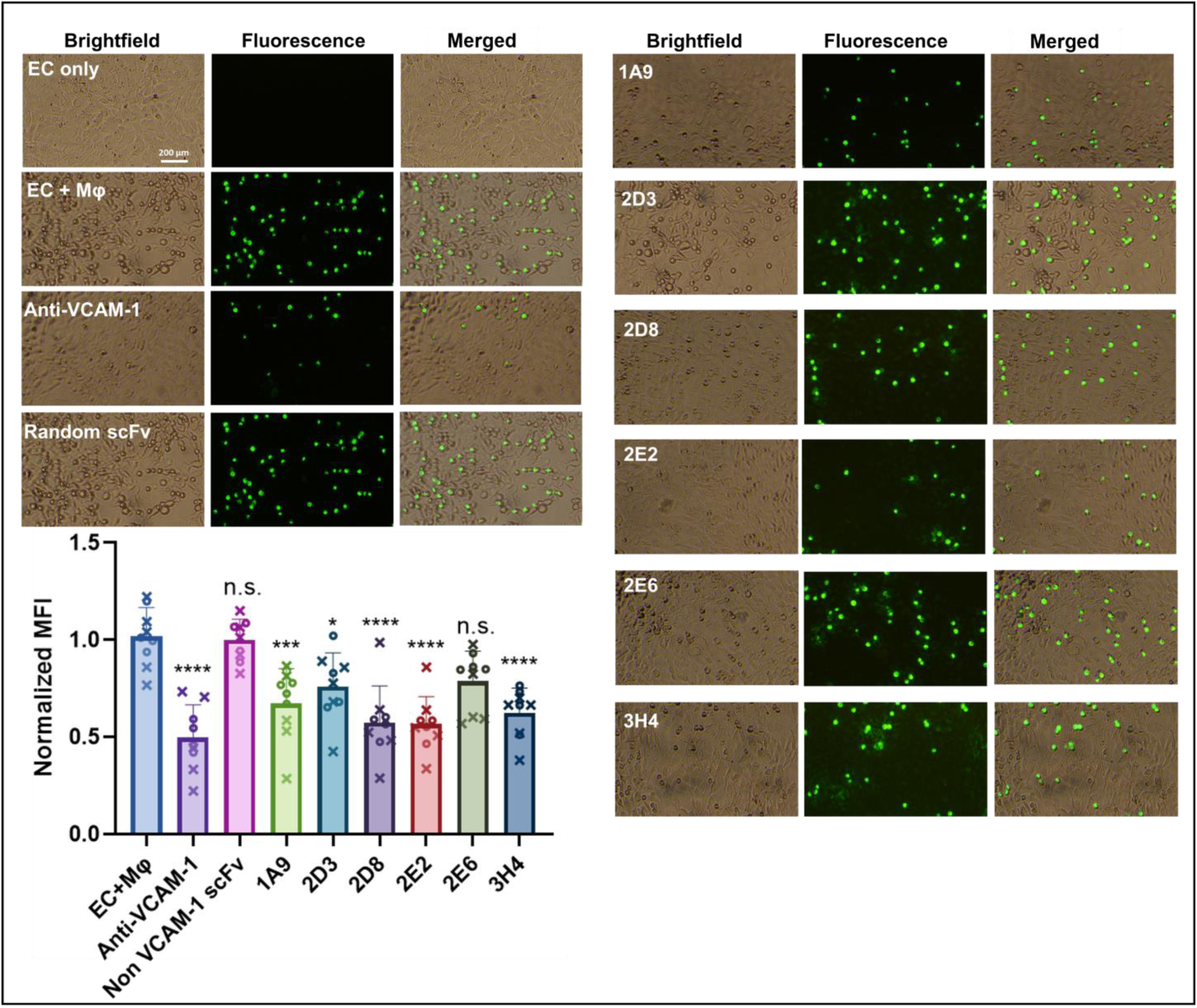
Anti-VCAM-1 scFvs targeting Ig-like domains 2 or 3 significantly decrease macrophage attachment to endothelial cells. Representative images from brightfield and green fluorescence, as well as their merged counterparts, show the attachability of macrophage J774 cells (green) to activated SVEC4-10 endothelial cells (EC) after treatment with our different antibodies and control antibodies (20 µg/ml). A non-antibody treatment and a random (not-VCAM-1 binder) scFv treatment were used as negative controls, while treatment with commercial full-length anti-mVCAM-1 (CBL1300) was used as positive control. Displayed is the average normalized MFI +/- SEM of two experiments with 4 or 5 replicates each, n = 9. The data was normalized by dividing each individual replicate to the mean of the non scFv treated control (EC+Macrophage) within each repeat. All statistical comparisons are to the EC+Macrophage control (One-Way ANOVA followed by post hoc Tukey pairwise comparisons): *p < 0.05, ***p < 0.001, ****p<0.0001; n.s. not significant.s

## 4. Discussion

In this project, we sought to identify antibodies that can bind mVCAM-1 at different Ig-like domains, which could be used to better dissect the biological function of different parts of VCAM-1 or for clinical applications. VCAM-1 is a cell surface glycoprotein implicated in the adhesion of white blood cells to endothelial cells during inflammatory processes. Due to VCAM-1’s role in a myriad of diseases, this molecule has been extensively studied and it has been targeted for therapeutic and diagnostic applications [1]. Multiple antibodies against VCAM-1 have been raised since this molecule was identified in the late 1980s ([51], **Supplementary Table S1**), and these have been in part used to study VCAM-1’s function. In some cases, both the binding site of these antibodies to VCAM-1 domains (especially to domains 1, 4, and 6) and the functional consequences of antibody binding are known, allowing a clear link between structure and function of VCAM-1 (**Supplementary Table S1**) [1, 8, 9, 19, 27–29, 32–34, 50–62]. However, in many other cases, only either the function or the potential binding site on VCAM-1 has been described, or neither information is available (**Supplementary Table S1**) [26, 60, 63–65]. In fact, the biochemical and biological role of most VCAM-1 domains remains poorly defined. In this study, we developed a suite of antibodies that bind to Ig-like domains 2, 3, or 5 of mouse VCAM-1 (with no apparent cross-reactivity with other VCAM-1 domains), which we hope will aid to more clearly dissect VCAM-1 molecular interactions, signal transduction, and physiological roles, and which may be used for downstream clinical applications.

The most highly expressed version of VCAM-1 in cells contains 7 Ig-like domains [8, 18, 19]. The function of Ig-like domains 1 and 4 has been thoroughly characterized, as they contain the binding site for one of VCAM-1’s most important binding partner, VLA-4 [1, 8, 9, 28]. Domain 2 appears to also be important for binding of α_1_β_4_ and, mainly, α_4_β_7_ integrins [29, 30]. In addition, VCAM-1 also mediates endothelial cell-eosinophil interaction [58, 62, 66], potentially through an interaction between eosinophils’ Galectin-3 [11, 13, 14] and VCAM-1’s Ig-like domains 3, 4, 5, and 6, which harbour sites for *N*-linked glycosylation. Interestingly, given *N*-linked glycosylation’s innate environment-dependent macro and microheterogeneity [67], the multiple potential sites for glycosylation in VCAM-1 could be powerful drivers of functional diversity in VCAM-1/Galectin-3 interactions. Domain 6, on the other hand, appears to be associated with TNFα-induced angiogenesis [31], and it is important for leukocyte transvasation but not adhesion [32]. Pepinsky et al [52] identified two protease-sensitive sites in 7D VCAM-1, one site between Ig-like domains 3 and 4 and one within two highly conserved cysteines in domain 5 (arguing for an unusual lack of disulfide bond formation in that Ig-like domain in VCAM-1). The functional implications of the biochemical characteristics of these two VCAM-1 regions are still unclear, but indicate differences in folding and accessibility to other proteins, including potential extracellular proteases that may regulate extracellular VCAM-1 shedding [52], internalization/degradation, and/or protein-protein interactions in a redox-sensitive manner. In addition, here we observed that deletion of domains 5, 6 and 7 together led to a reduction of cell surface expression of VCAM-1 (**Figure 4B, Supplementary Fig S6**). Even more, while the anti-VCAM-1 CBL1300 antibody bound well the Δ5, Δ7 and Δ6-7 mutants, this antibody bound the Δ5-7 mutant less efficiently (**Figure 4B, Supplementary Figure S6**), suggesting inappropriate folding of VCAM-1 or inaccessibility of the antibody to domain 1, which is the main binding site of CBL1300 (**Figure 4B, Supplementary Fig S6, and Supplementary Table S1**). A similar phenotype was observed with our 6 antibodies (**Figure 4C and Supplementary Figure S8**). We note that a secreted C-terminally Fc-tagged VCAM-1 construct carrying a Δ5-7 Ig-like domain deletion can be purified [32, 34]; however it is possible that the Fc C-terminal tag aided in this construct’s solubility and expression as a secreted molecule, which is different from our membrane bound construct. In fact, our results support Pepinsky *et al*.’s conclusion that VCAM-1 conformation depends on its C-terminal portion [52], and suggest a complex relationship among the last three Ig-like domains in determining expression and/or conformation of the full length VCAM-1 molecule. Altogether, our results and those of others argue for the need for a more detailed exploration of the biological roles of all VCAM-1 domains.

To identify new antibodies against VCAM-1 we performed phage display biopanning using a human naïve library, which is an animal-free platform for the discovery of monoclonal antibodies [40–42, 46, 68–70]. Biopanning is an affinity screening process which allows for the *in vitro* isolation of antibody-expressing phages from a phage display library that are specific to a target of interest [42, 70]. The target antigen was full length mouse 7D VCAM-1, a type-I transmembrane glycoprotein, and was presented through expression on the surface of mammalian cell lines, thereby ensuring its correct folding and post-translational modifications. Two rounds of biopanning were enough to generate an enriched pool of scFv-phages against mVCAM-1 (**Figure 1**). A third round of biopanning would have allowed us to further enrich the phage sub-library for anti-mVCAM-1 binders, but this may have also led to a reduction in the diversity of the scFv-phage population by selecting against weaker or more poorly expressed binders that may bind different parts of mVCAM-1. We identified several antibodies that bound mVCAM-1 as scFv-phage (**Figure 2**), as scFv (**Figure 3**), and as full length mAb (**Figure 4**). Two of the scFv-phage binders were not pursued, due to the presence of sequences in their CDRs that could lead to potential folding/binding errors (clones 1H3 and 2B9) when expressed in bacteria and/or mammalian cells (data not shown). Three of the 6 antibodies identified bound mVCAM-1 Ig-like domain 3: 1A9, 2D8, and 2E2; two bound Ig-like domain 2: 2D3 and 3H4; and one bound Ig-like domain 5: 2E6 (**Figure 4**). Interestingly, all antibodies reduced macrophage attachment to endothelial cells when compared to the non-antibody treated control (EC+macrophage), with 5 of them significantly reducing macrophage attachment: clones 1A9, 2D3, 2D8, 2E2, and 3H4, which are Ig-like domain 2 or 3 binders (**Figure 5**, p < 0.05, **Supplementary Figure S8**; we note that clone 2E6 showed p = 0.056). Similar results were obtained when the full length IgG2a versions of these antibodies were tested in the same assay, except that all antibodies, including clone 2E6, significantly reduced macrophage attachment to endothelial cells (Pickett et al., manuscript in preparation). Different VCAM-1 binding antibodies will require different concentrations to exert a functional effect due to different affinities for the antigen (a property of the paratope/epitope sequence and of accessibility to the epitope). An antibody titration experiment would allow for a more detailed comparison of the different antibodies. Overall, our results show that blocking other VCAM-1 Ig-like domains beyond 1 and 4 [1, 9, 21, 28] can also reduce cell-to-cell attachment (**Figures 4 and 5**).

There are multiple ways in which anti-VCAM-1 antibodies could impact VCAM-1 mediated attachment and transmigration. Hession *et al*. [19] hypothesized early on that VCAM-1 may play other roles beyond being an adhesin, including participating on signal transduction. Indeed, VCAM-1 activation leads to cytoskeleton remodeling and changes at the endothelial cell junctions [71, 72], and antibodies against VCAM-1 can affect VCAM-1 downstream signaling [17, 34, 72]. Some of the ways antibodies can alter VCAM-1 function is by directly or indirectly blocking protein-protein interactions or by altering VCAM-1 downstream signaling in a ligand-independent manner. To do this, antibodies could block access to residues involved in the interaction, or alter VCAM-1 epitope accessibility, folding/conformation, quaternary structure (homo or hetero multimerization), or cell surface expression by e.g. triggering VCAM-1 internalization. For example, our Ig-like domain 2 binders could be blocking attachment by preventing proper interaction with α_4_β_7_ integrin [30]. Additional experiments using molecular docking, mutagenesis, and structural biology to dissect antibody-VCAM-1 interactions, a more detailed study of the precise VCAM-1 epitope to which each antibody is binding, and the functional impact of changes on/blocking of these domains on protein-protein interactions, cellular physiology, and downstream signaling will help in understanding the structure-function relationship of each VCAM-1 Ig-like domain and how antibodies impact them. Given that scFvs are considerably smaller than full length monoclonal antibodies (∼25 kDa *vs* ∼150 kDa, respectively), the chances that the impact of a monomeric scFv on attachment is due to steric hindrance of the VLA-4/VCAM-1 interaction is low (although this would have to be demonstrated biochemically, which lies outside the scope of this manuscript). Thus, the antibodies described in this work could help uncover new VCAM-1 biology (binding partners and/or function).

Gaining a deeper knowledge of VCAM-1 functions would not only facilitate our general understanding of physiology and biology, but it would also provide key information for the design of precision therapies targeting specific structural domains associated with specific clinical outcomes. Competition studies with purified VLA-4 or other known binders and antibodies would help elucidate potential novel mechanisms of action. In addition, researchers could harness the power of omic techniques and AI to perform a more comprehensive spatial and temporal interactomic, transcriptomic, proteomic, phospho-proteomic, and/or glycoproteomic profiling of VCAM-1 (WT and Ig-like domain mutants) and downstream signaling pathways, in the presence or not of ligands and/or anti-VCAM-1 antibodies. These analyses could help uncover signaling pathways, new interacting proteins, and the function of specific Ig-like domains. Analyses targeting post-translational modifications in VCAM-1 or its ligands are especially important to understand how these modifications impact VCAM-1 function in health and disease (e.g. during cancer or inflammation). These antibodies could also help better dissect the role of VCAM-1 amino acid sequences involved in ligand interaction [30]. Despite the large amount of data already acquired on VCAM-1, there is still plenty to learn about this molecule’s physiological roles.

Antibodies against VCAM-1 could have a range of clinical applications in cardiovascular disease, cancer, and inflammatory diseases. For instance, it would be interesting to test these antibodies in preclinical animal models of atherosclerosis. The antibodies (either in IgG or scFv formats) can also be used as antibody-drug conjugates, to specifically deliver payloads with anti-cancer or anti-inflammatory properties. However, further studies need to be conducted to determine whether binders also exert an activating function, since activating VCAM-1 downstream signaling may be counterproductive to the desired clinical outcome. Anti-VCAM-1 antibodies could also be used for diagnostic purposes, to identify VCAM-1 expressing cancer or atherosclerotic plaques, etc. Cross-reactivity of these antibodies with other Ig-like domain containing proteins could be a potential issue that may require further *in silico* and *in vitro* testing before moving into pre-clinical models. Further progression into clinical studies would require the identification of cross-reacting antibodies to both mouse and human VCAM-1. Anti-VCAM-1 antibodies could have a range of therapeutic, diagnostic, or theranostic applications.

For the purpose of the functional assays, this work focused on scFv anti-VCAM-1 versions, rather than the full-length antibodies for two main reasons. First, if an impact on attachment was observed with the scFv, this impact would highly likely also be observed with the full-length antibody, while the reverse is not necessarily true. Second, scFvs have many advantages over full length antibodies in terms of biochemistry and pharmacokinetics [73, 74]. A typical mammalian IgG antibody is a large molecule consisting of two heavy chains linked to each other and to a light chain by disulphide bonds [38, 73–75]. On the other hand, a scFv is a small functional antigen binding domain that consists of one variable heavy and one variable light domain joined by a peptide linker, ∼ 1/6^th^ the size of a full length IgG [73, 74, 76]. Compared to full length antibodies, the smaller size of scFvs allows for better tissue penetration and rapid clearance while maintaining target binding specificity [73, 74, 77]. The enhanced tissue penetration is especially advantageous in some pathologies such as cancer [78] and atherosclerosis [35], in which scFvs can reach the tumour or plaque core better than full length antibodies [73, 74, 77, 79]. Further, rapid clearance is useful when antibodies are carrying a toxic payload (e.g. for radioactive immunotherapy or diagnostics) [80]. The scFvs also have reduced immunogenicity and no effector function, since they lack the Fc region [81]. There are many differences between scFv and full-length antibodies, and the decision on which kind of antibody to use will ultimately depend on the specific biochemical test required or desired therapeutic modality.

Here, we showed that scFvs targeting different Ig-like domains of VCAM-1 that are not directly associated with the VLA-4/VCAM-1 interaction can also negatively impact macrophage-endothelial cell attachment (**Figures 4 and 5**). Our data emphasizes that there is still plenty to learn about VCAM-1 biological function, and our antibodies are excellent tools to facilitate future studies. Given the critical role this molecule plays in a myriad of physiological and disease states, understanding its biochemical characteristics and molecular interactions will be key to designing more precise and targeted future therapies.

## Author Contributions

Conceptualization, Lucia Zacchi; Data curation, Binura Perera and Lucia Zacchi; Formal analysis, Binura Perera, Yuao Wu and Lucia Zacchi; Funding acquisition, Martina Jones, Hang Ta and Lucia Zacchi; Investigation, Binura Perera, Jessica Pickett, Yuao Wu, Francisca Barretto and Lucia Zacchi; Methodology, Yuao Wu, Nadya Panagides, Christian Fercher, David Sester, Martina Jones, Hang Ta and Lucia Zacchi; Project administration, Lucia Zacchi; Resources, Martina Jones, Hang Ta and Lucia Zacchi; Supervision, Hang Ta and Lucia Zacchi; Writing – original draft, Binura Perera, Yuao Wu, Francisca Barretto and Lucia Zacchi; Writing – review & editing, Binura Perera, Christian Fercher, David Sester, Martina Jones, Hang Ta and Lucia Zacchi. All authors have read and agreed to the published version of the manuscript.

## Data Availability Statement

The raw data supporting the conclusions of this article will be made available by the authors on request. Antibody sequences will not be made available to protect commercialization potential.

## Acknowledgments

We thank Dr Zhao Wang from the Rowan lab at The University of Queensland for the confocal microscopy images, and Dr Christopher Howard for his help with protein purification.

