## Supplementary Information for "Isolation and Characterization of Antibodies Against VCAM-1 Reveals Putative Role for Ig-like Domains 2 and 3 in Cell-to-Cell Interaction"

*General information on phage display biopanning*

Phage display antibody biopanning requires the generation of an antibody library fused to a phage coat protein that is expressed on the surface of the phage [1, 2]. To construct a phage display library, the gene fragments encoding the variable regions of the heavy and light chains of antibodies are extracted from the mRNA of B cells from "healthy/naïve" (no disease/immunization) or "immune" (sick/vaccinated/immunized) donors. These variable regions are assembled into small antigen binding units such as scFvs, and are fused to the phage G3P coat protein for phage surface expression, thereby generating a phage display library in which each phage expresses a unique scFv-G3P fusion [3, 4]. These libraries are used for biopanning against a specific antigen [5]. Biopanning consists of four basic steps: scFv-phage binding to antigen, washing to reduce unspecific phage binding, elution of bound scFv-phages, and infection of bacteria with eluted bound phages to generate enriched sub-libraries [6]. This cycle is repeated for 2-4 rounds until specific binders are enriched and identified [2, 6-9].

### Supplementary methods

*mVCAM-1 DNA sequence for mVCAM-1-eGFP vector design*

The DNA sequence encoding the open reading frame of mouse VCAM-1 (Uniprot ID P29533) was codon optimized for expression in *Cricetulus griseus* (CHO cells). A Kozak sequence, and NheI, BamHI, and EcoRI restriction sites were added to the 5' and 3' ends for cloning purposes, as shown below.

**Optimized sequence with NheI and KOZAK at the 5' end and BamHI-stop-EcoRI at the 3' end:**
**GCTAGC**cacc
ATGCCAGTGAAGATGGTGGCAGTGCTAGGTGCTTCCACCGTCCTTTGGATTCTGTTCGCC
GTTTCCCAGGCCTTCAAGATCGAGATCAGCCCTGAGTACAAGACCATAGCCCAGATCGGC
GATTCCATGGCCCTGACCTGCTCTACAACCGGCTGTGAATCTCCACTGTTTTCCTGGCGG
ACACAGATCGACTCCCCACTGAACGCCAAGGTGAGAACCGAGGGCTCTAAGTCCGTGCTG
ACCATGGAACCTGTGTCCTTCGAGAACGAACACTCTTACCTGTGCACCGCTACCTGCGGC
TCCGGCAAACTGGAGCGCTCCATCCACGTGGACATCTACTCATTTCCCAAGGACCCTGAG
ATCCAGTTCTCTGGCCCTCTGGAAGTGGGGAAGCCTGTGACAGTGAAGTGCCTGGCTCCT
GACATCTACCCTGTGTACAGACTGGAAATCGACCTGTTCAAGGGCGATCAGCTGATGAAC
AGACAAGAGTTCTCCTCTGAAGAGATGACCAAGTCTCTGGAGACAAAGTCTTTGGAGGTC
ACCTTCACCCCCGTGATCGAAGACATCGGAAAGGCTCTGGTGTGCAGAGCCAAACTGCAC
ATCGACCAGATCGATTCTACACTCAAAGAGAGAGAGACAGTGAAGGAGCTCCAAGTGTAC
ATCTCCCCTAGAAACACCACTATCTCTGTGCATCCTTCCACCAGACTGCAAGAGGGCGGC
GCCGTGACCATGACCTGCTCCTCTGAGGGCCTGCCTGCCCCTGAGATCTTCTGGGGACGG
AAGCTGGACAACGAGGTGCTGCAGCTGCTGTCTGGAAATGCCACCCTGACACTGATCGCC
ATGCGGATGGAAGATTCCGGCGTGTACGTGTGCGAGGGAGTGAACCTGATCGGCCGGGAC
AAGGCCGAGGTGGAACTGGTCGTGCAGGAAAAGCCCTTCATCGTGGATATTTCCCCTGGC
AGCCAGGTGGCTGCTCAGGTGGGAGATTCCGTGGTGTTGACATGTGCTGCCATCGGCTGC

GACTCCCCCTCTTTCTCCTGGCGGACCCAGACCGACTCGCCTCTGAACGGCGTAGTGAGA
AACGAGGGCGCTAAGTCTACCCTCGTGCTGTCTTCTGTCGGCTTCGAGGACGAGCATTCT
TACCTGTGTGCTGTGACCTGCCTGCAGAGAACCCTGGAGAAGCGGACCCAGGTGGAAGTG
TACAGCTTCCCTGAGGACCCTGTAATCAAGATGTCCGGACCTCTGGTCCACGGCAGACCC
GTGACCGTTAACTGCACCGTGCCCAACGTCTATCCTTTCGACCACCTGGAGATCGAGCTG
CTGAAGGGCGAAACCACACTGATGAAGAAGTACTTCCTGGAAGAAATGGGCATCAAGAGC
CTTGAGACCAAGATCCTGGAAACCACCTTTATCCCTACAATCGAGGATACCGGCAAGTCC
CTGGTGTGTCTGGCCCGGCTGCACTCCGGCGAGATGGAATCCGAGCCTAAGCAGCGGCAG
TCCGTGCAGCCTCTGTACGTGAACGTGGCCCCTAAGGAAACCACCATCTGGGTGTCCCCT
AGTCCCATCCTGGAGGAAGGCTCCCCAGTGAATCTGACCTGTTCTTCCGATGGCATCCCA
GCGCCTAAGATCCTGTGGTCCAGACAGCTGAACAACGGCGAGCTGCAGCCCCTGAGCGAG
AATACCACCCTGACCTTCATGAGCACCAAGAGAGACGACTCTGGAATCTATGTGTGCGAA
GGCATCAACGAGGCCGGCATCAGCCGGAAGTCCGTGGAACTGATCATCCAAGTGTCTCCT
AAAGACATCCAGCTGACCGTGTTCCCCTCCAAGTCCGTGAAAGAGGGCGACACCGTGATC
ATCTCCTGCACCTGTGGCAACGTGCCTGAGACCTGGATCATCCTGAAGAAGAAAGCTAAG
ACAGGCGACATGGTCCTGAAGTCCGTGGACGGCTCCTACACCATCAGACAGGCCCAGCTG
CAGGACGCTGGCATCTACGAGTGCGAGTCTAAGACCGAGGTGGGCTCTCAGCTCCGCTCC
CTGACATTGGACGTGAAGGGGAAAGAGCACAACAAGGACTACTTCTCTCCTGAGCTGCTG
GCTCTGTACTGCGCCAGCTCCCTGGTGATCCCTGCCATCGGCATGATCGTGTACTTTGCC
AGAAAGGCCAACATGAAGGGCTCCTACTCTCTGGTGGAAGCTCAGAAATCTAAAGT
GGATCCatgagaattc

*Generating the scFv-LPETG versions via site-directed mutagenesis*

To generate a pET28b_LPETG vector, the following sequence ttaaCCATGGGATATCAAGCTTGCGGCCGC**CTGCCAGAAACCGGT**CTCGAGttaa was added to pET28b (equivalent to ttaa-NcoI-EcoRV-*HindIII*-NotI-**LPETG**-XhoI-ttaa). To do this, two oligomers covering this entire sequence complementary to each other (Sigma Aldrich) were resuspended to a final concentration of 100 mM and annealed together in a thermocycler. Annealing was achieved by heating the sample to 95 °C for 2 minutes and slowly cooling down to 25°C, over a span of 45 minutes. The annealed oligomers were then digested with the restriction enzymes NcoI and XhoI (New England BioLabs® Inc.). An empty pET28b vector was digested simultaneously with the same enzymes, dephophorylated with alkaline phosphatase, and purified using the Qiagen Qiaquick Gel Extraction Kit. Following the digestion of the annealed oligos, the enzymes were heat inactivated at 80 °C for 20 minutes before ligating the annealed oligo with the purified, linearized pET28b vector. The ligation was incubated at 16 ˚C overnight followed by heat inactivation of the enzymes at 65 °C for 10 minutes and then the reaction was held at 4 °C in the Veriti™ 96-Well Thermal Cycler (Applied Biosystems™, 4375786). The ligation was transformed into competent *E. coli* DH5a cells following standard protocols to generate pET28b-LPETG*. The next day the colonies were tested by colony PCR and verified by sequencing, as described in section 3.6. A frameshift was inadvertently incorporated in the sequence during the design. To correct this frameshift, site-directed mutagenesis was performed using the following primers, LPETG codon corrected FWD (ACTGCCAGAAACCGGTCTCG) and LPETG codon corrected REV (GCGGCCGCAAGCTTGATATCC) (**Supplementary Table S3**). The binding of the FWD primer introduces an overhang of 1 base pair which will be amplified with the pET28b-LPETG* plasmid. The Q5® High-Fidelity DNA Polymerase was used to amplify the vector following manufacturer's recommendations and considering the primers annealing temperature. The PCR product was purified using the QIAquick® PCR Purification Kit (Qiagen) and the linearized DNA was phosphorylated by T4 PNK (New England BioLabs® Inc.) using 100 ng of the purified linear DNA in a reaction mixture containing 5 mL of T4 PNK Reaction Buffer (10X), 5 mL of ATP (10 mM), 1 mL of T4 PNK and nuclease free water to a final volume of 50 mL. The reaction was incubated for 30 minutes at 37 °C and the enzymes were heat inactivated at 65 °C for 20 minutes. The phosphorylated DNA was then ligated using T4 ligase and the ligation transformed into competent *E. coli* DH5a cells to generate plasmid pET28b-LPETG (LZCBI53). Next, the DNA encoding the selected scFvs was subcloned from the phagemid vectors into pET28b(+)-LPETG (LZCBI53) vector through NcoI and NotI double restriction digestion and ligation (Supplementary Table S2). All vectors were sequenced.

*mVCAM-1 mutagenesis strategy to generate Ig-like domain mutants*

The deletion of mVCAM-1 Ig-like domain(s) starts and ends in the middle of the inter-domain sequence immediately before or immediately after the domain(s) deleted. All mutagenesis procedures were performed using Q5 High-Fidelity DNA Polymerase (NEB, M0491) and primers described in **Supplementary Table S3**. To make these constructs, an N-terminal FLAG tag was first introduced after the signal sequence into LZCBI18 to generate pEGFP-N1_mVCAM-1(1-24)-FLAG-mVCAM1(25-793) (LZCBI120) using primers mVCAM-1 FLAG Fwd and mVCAM-1-FLAG Rev (**Supplementary Table S3**). All Ig-like deletions were made on the LZCBI120 backbone, and 9 different constructs were made: ΔIg-like-1 (primers mVCAM1-d1 Fwd and mVCAM1-d1 Rev), ΔIg-like-1-2 (primers mVCAM1-d2 Fwd and mVCAM1-d1 Rev), ΔIg-like-1-3 (primers mVCAM1-d3 Fwd and mVCAM1-d1 Rev), ΔIg-like-4-7 (primers mVCAM1-d7 Fwd and mVCAM1-d4 Rev), ΔIg-like-5-7 (primers mVCAM1-d7 Fwd and mVCAM1-d5 Rev), ΔIg-like-6-7 (primers mVCAM1-d7 Fwd and mVCAM1-d6 Rev), ΔIg-like-7 (primers mVCAM1-d7 Fwd and mVCAM1-d7 Rev), ΔIg-like-3 (primers mVCAM1-d3 Fwd and mVCAM1-d3 Rev), and ΔIg-like-5 (primers mVCAM1-d5 Fwd and mVCAM1-d5 Rev) (**Supplementary Table S3**). PCR amplified DNA fragments were cleaned with Nucleospin Gel and PCR clean-up mini kit (Macherey-Nagel, 740609), phosphorylated with T4 polynucleotide kinase (NEB, M0201), ligated with T4 Ligase (NEB, M0202), and transformed into chemical competent *Escherichia coli* DH5alpha strains or ST0213 Stellar competent cells (Takara, 636766).

### Supplementary Figures

*Expression of mVCAM-1_GFP*

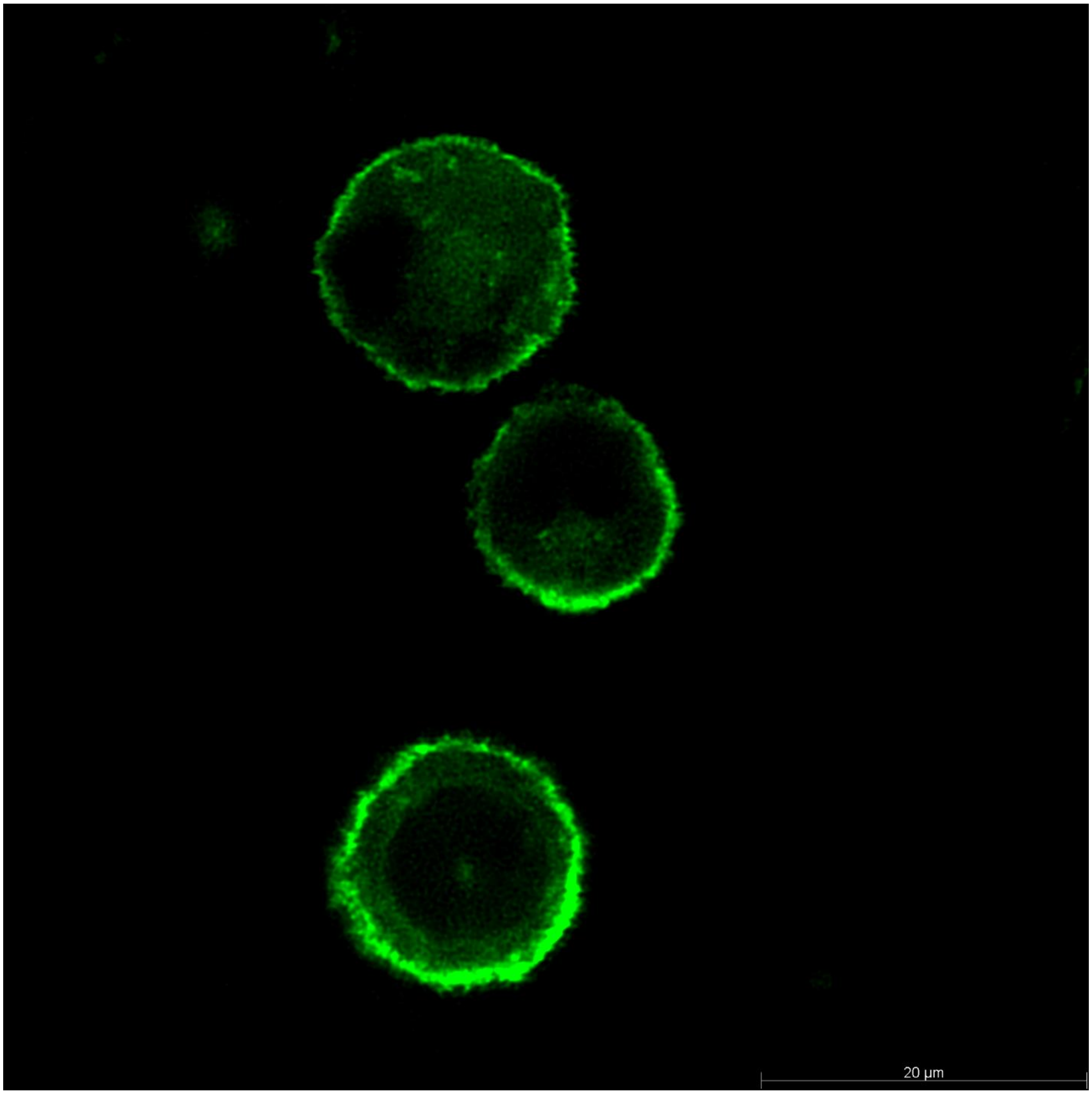


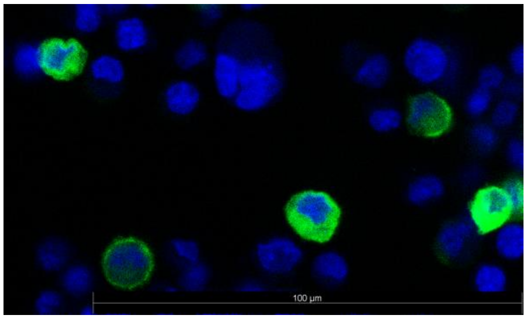

**Figure S1**. **mVCAM-1-eGFP localizes to the cell periphery in mammalian cells.** Confocal microscopy of CHO XL99 cells transfected with pEGFPN-1_mVCAM-1 showed that mVCAM-1-eGFP is expressed and localized at the cell periphery. The DAPI+GFP overlay image also shows non-expressing cells, demonstrating that the green fluorescence observed is indeed mVCAM-1-eGFP, and not autofluorescence of the cells. Images courtesy of Zhao Wang, Rowan Lab, The University of Queensland.

*Biopanning –flow cytometry of phage pools*

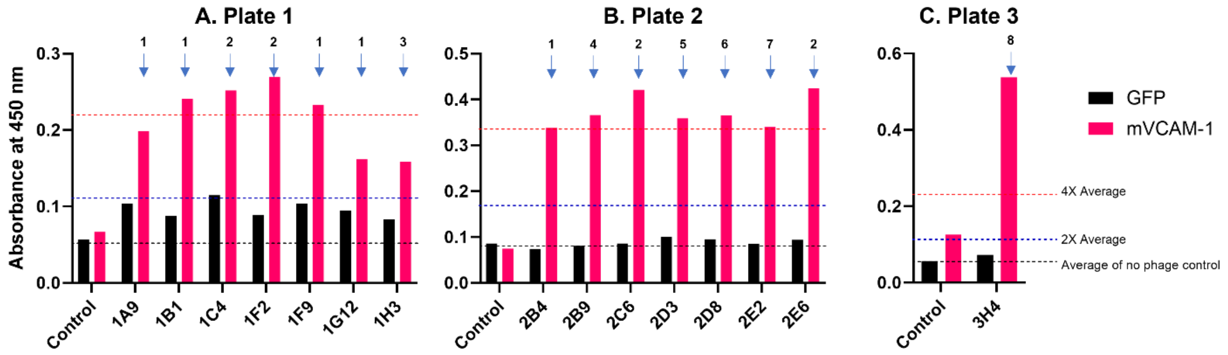


**Supplementary Figure S2. Monoclonal whole cell ELISA of round 2 phage pool showed 15 positive clones**. Three 96-well plates with individual clones from round 2 phage pool were screened, and the 15 positive clones identified from Plate 1 (**A**), Plate 2 (**B**), and Plate 3 (**C**) are shown. The black line in the graphs indicates the average of the no phage controls in each plate, and the blue and red lines indicate 2-times and 4-times the value of this average, respectively. There are two negative controls in this experiment. First, cells expressing eGFP or mVCAM-1-eGFP incubated only with primary and secondary antibodies but with no phages. The absorbance value of these no-phage controls determined the value of the black line: average of no-phage controls (black line). Second, cells expressing eGFP incubated with phages. When testing the specific isolated phage clones, our criteria to select a positive clone required that the absorbance of a particular phage clone on the eGFP negative control would not exceed 2-times the value of the no phage controls (blue line), and that the corresponding absorbance on the mVCAM-1-eGFP expressing cells was equal or above 4-times the value of the no phage controls (red line). The reason we allowed higher absorbance on the eGFP-only cells above the no phage control is to account for the possibility that inefficient washes or phage stickiness eliminate perfectly good candidates as false negatives. The 4-times above the average requirement to determine a true positive increased the stringency for selection, thereby reducing the chances of selecting false positives. Many of the recovered clones shared the same DNA sequence. All unique positive sequences are listed with a number on the graph, with positive clones with identical sequences sharing the same number.

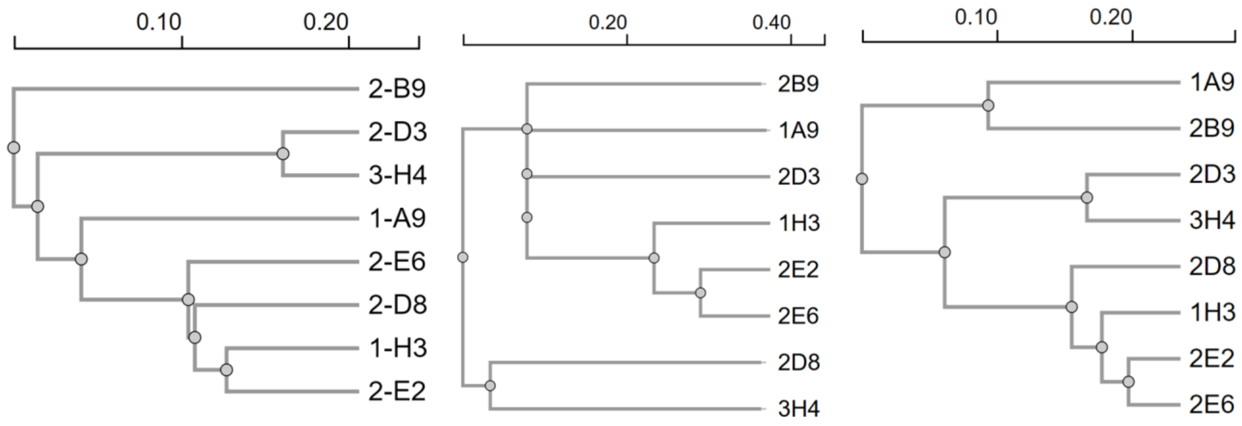


**Supplementary Figure S3. Phylogenetic tree of the 8 unique scFv sequences identified.** A phylogenetic tree was generated using the entire amino acid sequence of the 8 unique scFvs identified, using Clustal Omega, to facilitate comparing similarities among sequences. The alignment on the left shows the phylogenetic tree for the entire scFv sequences, which reflects differences and similarities in all CDRs and frameworks, while the one on the middle is only for the CDR3 of heavy chains, and the one on the right is only for the entire light chains. As observed on the left and on the right, clones 2D3 and 3H4 are very similar overall and, coincidentally, they also bind the same Ig-like domain in mVCAM-1. As observed on the right, 1A9 and 2B9 are clearly separated from the other antibodies since these two contain kappa light chains while the others contain lambda light chains. Also as observed, on the right, clones 2E2 and 2E6 cluster closely together as they share an almost identical light chain (even though they bind two different Ig-like domains, suggesting a higher role for the heavy chain in determining the epitope).

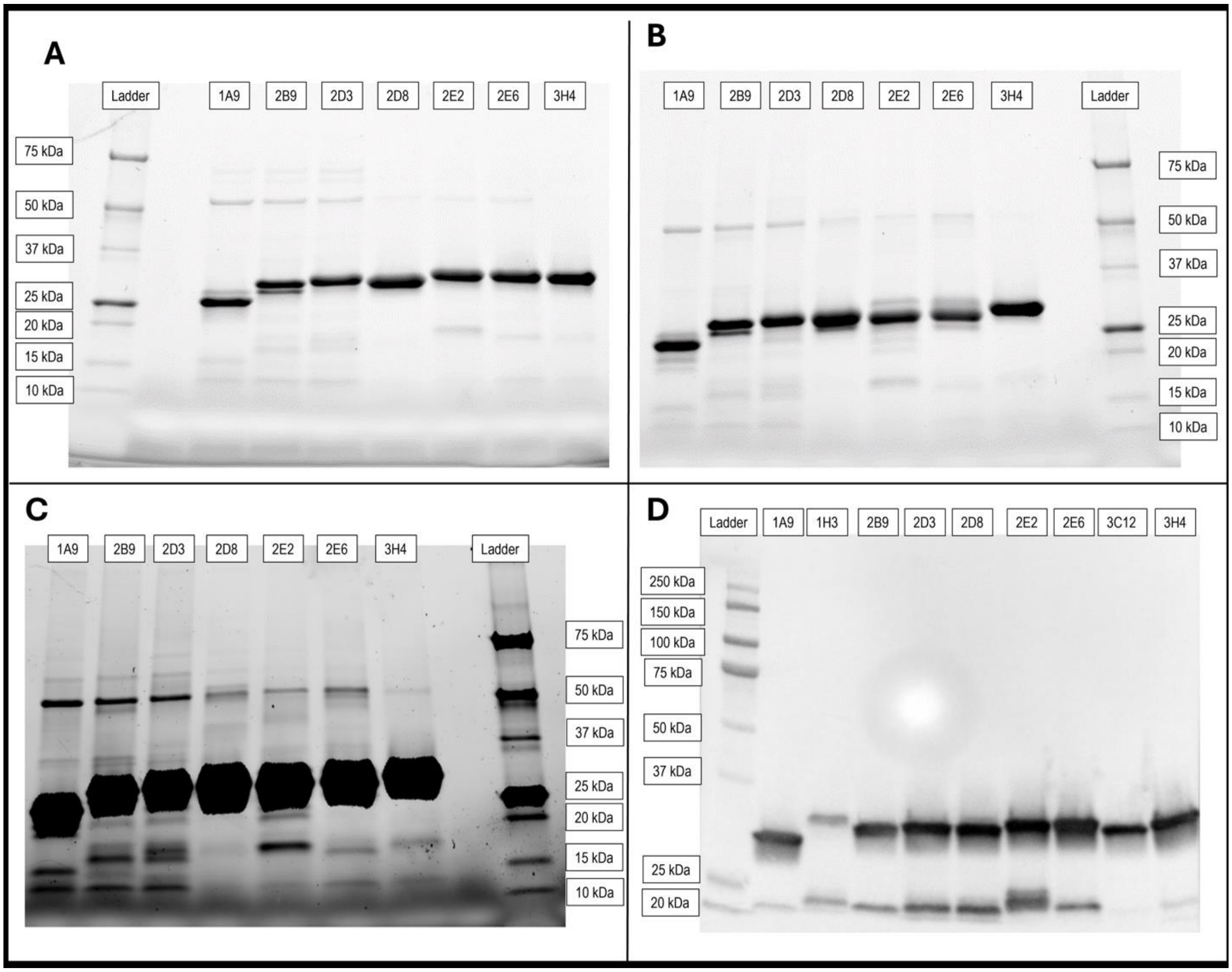


**Supplementary Figure S4**. **SDS-PAGE of purified scFv-LPETG clones, and western blot of cell extracts pre-purification.** Reducing **(A)** or non-reducing **(B,C)** SDS-PAGE of purified scFv-LPETGs. Panel (**C**) is an overexposed version of the gel in **(B)**. The gels were loaded with 5 µg of purified scFv-LPETG of clones 1A9, 2B9, 2D3, 2D8, 2E2, 2E6, and 3H4. **D)** Western blot of whole cell extracts from SHuffle *E. coli* strains expressing the different scFv-LPETGs prior to purification. The blot was prepared as described in the methods in the main text. The anti-His-HRP (Miltenyi Biotec Cat# 130-092-785, RRID:AB_1103231) antibody used was at a 1:5,000 dilution.

We note that this figure contains the clones 1H3 (**D**), 2B9 (**A-D**), and 3C12 (**D**). As described in the main text, clones 1H3 and 2B9 were not further pursued due to potential sequence liabilities when expressed in eukaryotic cells, as explained in the results section. Clones 2B9 and 3C12 bound to mVCAM-1 expressing cells with high specificity, while clone 1H3 showed extensive background binding (data not shown). Clone 3C12 has not been discussed in this manuscript.

The antibodies were purified with varied levels of purity, and additional proteins can be seen co-purifying with the antibodies with different relative abundances. Mass spectrometry analysis of one of the purified scFv samples (clone 2D8) showed that the 3 most abundant co-purified proteins had MW in the range of 55-70 kDa (see **Supplementary Table S5**). Bands with similar MW to these three proteins were generally observed in all purified scFvs and scFvs-LPETG clones

(**Supplementary Figure S4** and data not shown). We observed similar behaviour of the scFv-LPETG tagged antibodies compared to the scFv antibodies in the SDS-PAGE from purified samples, western blot from cell extracts pre-purification, and by flow cytometry (data not shown) indicating that, as expected, the tag is unlikely to have an impact on antibody expression and function.

We note that in the western blot (**Supplementary Figure S4D**) there is an ~15 kDa or lower MW band reactive with anti-HIS antibody observed in all the cell extracts, with different intensity levels. These bands could be partially cleaved antibodies that still contain the C-terminus. However, these lower MW fragments are either not observed in the purified material or present at considerably lower relative abundances (compare D with B,C), possibly because the fragments were lost during the dialysis step. Therefore, these lower MW fragments (containing the light variable chain) are unlikely to have had any impact on the subsequent functional analysis performed on these antibodies.

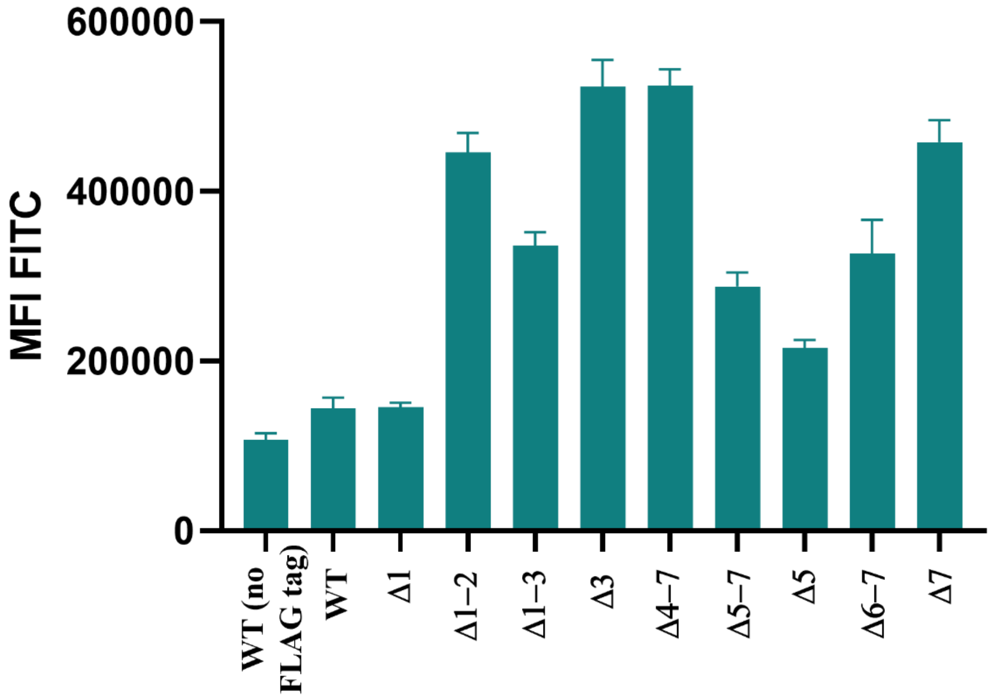


**Supplementary Figure S5**. **Flow cytometry data comparing GFP expression for all Ig-like domain mutants.** ExpiCHO cells transiently transfected with mVCAM-1-eGFP WT (with or without N-terminal FLAG) or FLAG-tagged mVCAM-1-eGFP Ig-like deletion constructs were used for flow cytometry as described in the methods. Overall expression efficiency of the constructs was determined using GFP expression (FITC signal). Results represent the average MFI (FITC-H) +/- SE from one experiment with 21 replicates for each vector. The data shows that addition of the FLAG tag or deletion of Ig-like domain 1 only marginally increased the overall GFP expression in transfected cells. However, all other constructs showed a larger increase in GFP signal that did not always correlate with anti-FLAG antibody binding (a marker for cell surface expression, **Figure 4**, **Supplementary Figure S6**) and could indicate intracellular accumulation of GFP (potentially due to folding defects). Importantly, the Δ5-7 Ig-like mutant construct (which showed poor binding to all antibodies tested, **Figure 4, Supplementary Figure S6**) showed higher overall GFP expression than the controls WT mVCAM-1 with or without N-terminal FLAG tag), demonstrating that lack of antibody binding to this construct by flow cytometry was not due to lower overall construct expression.

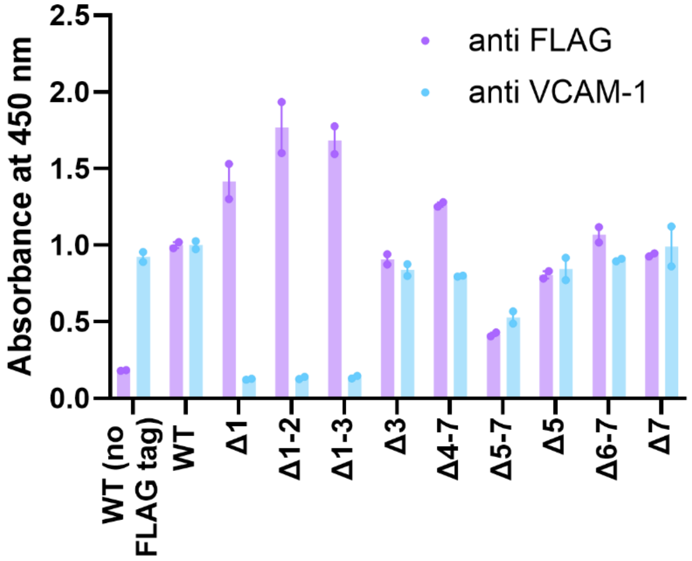


**Supplementary Figure S6**. **Whole cell ELISA comparing cell surface expression for mVCAM-1-eGFP constructs.** ExpiCHO cells transiently transfected with mVCAM-1-eGFP WT (with or without N-terminal FLAG) or FLAG-tagged mVCAM-1-eGFP Ig-like deletion constructs were used for whole cell ELISA. In this experiment, $5x10^5$ cells/well were stained with either 1:400 rabbit anti-FLAG (Cell Signaling Technology Cat# 2368, RRID:AB_2217020) or 1:400 rat anti-VCAM-1 CBL1300 (Millipore Cat# CBL1300, RRID:AB_2214062) primary antibodies, followed by 1:1,000 anti-rabbit IgG-HRP (Cell Signaling Technology Cat# 7074, RRID:AB_2099233) or 1:1,000 anti-rat IgG-HRP (Cell Signaling Technology Cat# 7077, RRID:AB_10694715) secondary antibodies. Graphs show the average-Max-Min of the Absorbance at 450 nm from one experiment with 2 replicates for each vector. All results are normalized to the values of the mVCAM-1-eGFP-FLAG tagged WT vector. The results of this experiment are in complete agreement with the results from the flow cytometry experiment (**Figure 4**), and support that: 1) addition of the FLAG tag did not lead to a decrease in cell surface expression of the mutant mVCAM-1 versions; 2) CBL1300 binds mVCAM-1 Ig-like domain 1 (and **Supplementary Table S1**); and 3) cell surface expression of the Δ5-7 Ig-like mutant construct is less efficient compared to the other constructs (the differences between the flow cytometry data in **Figure 4** and the ELISA data in this figure are likely due to differences in the sensitivity of the assays in these specific experimental conditions, and support that this construct is indeed at least partially expressed on the cell surface (see discussion)).

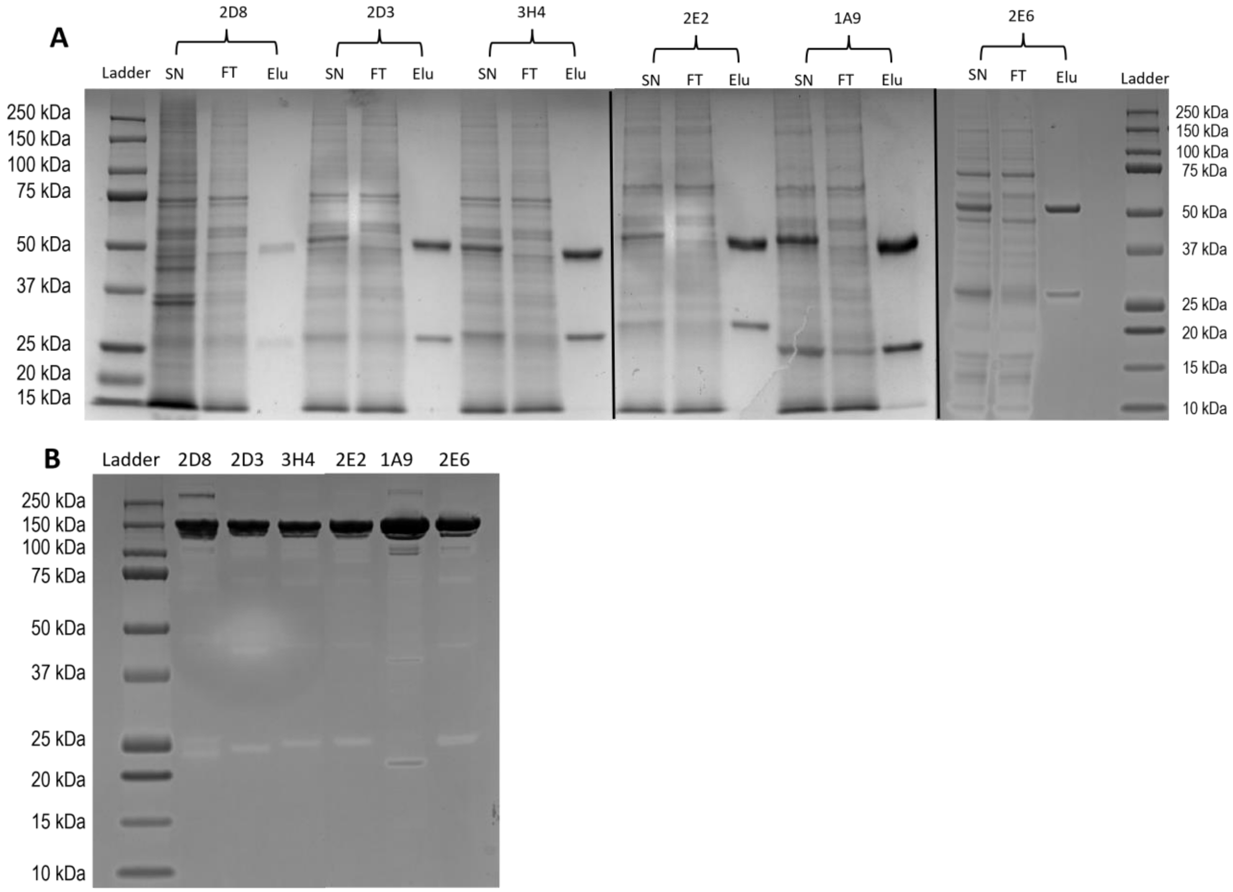


**Supplementary Figure S7. Reducing and non-reducing SDS-PAGEs of full-length reformatted IgG2a monoclonal anti-mVCAM-1 antibodies. A)** Reducing SDS-PAGE of the supernatant (SN), Flow-through (FT), and 5 µg of purified mAb (Elu) for clones 1A9, 2D3, 2D8, 2E2, 2E6, and 3H4. **B)** Non-reducing SDS-PAGE showing 5 µg of purified mAb for clones 1A9, 2D3, 2D8, 2E2, 2E6, and 3H4.

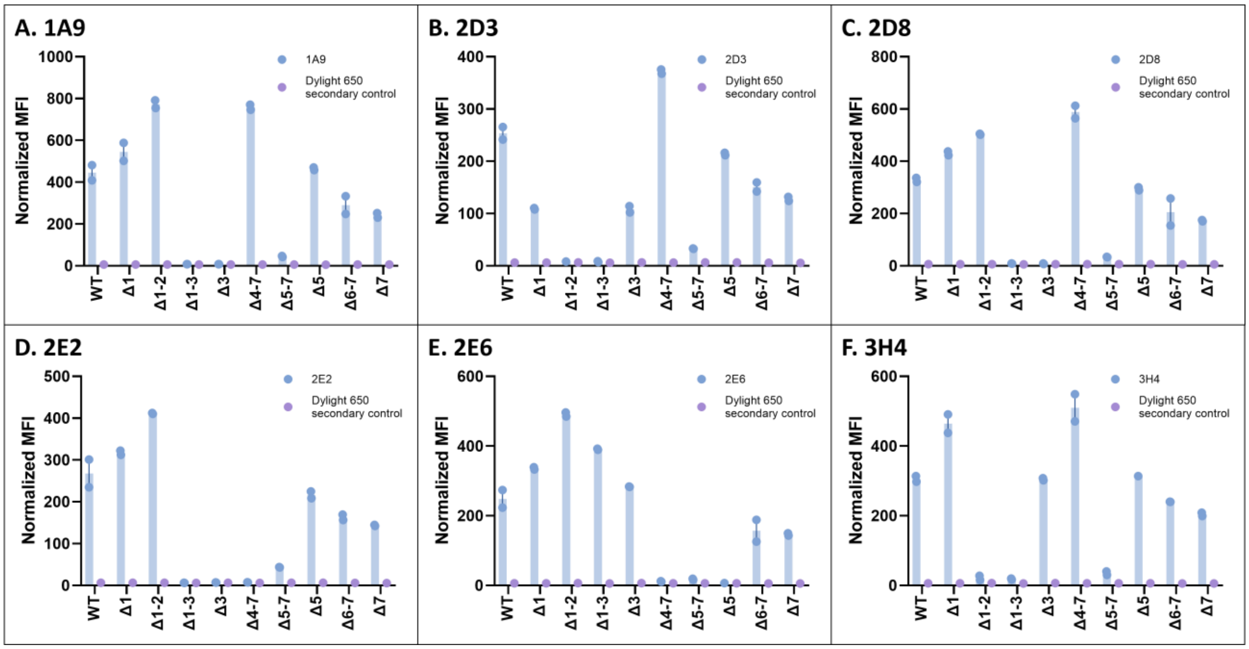


**Supplementary Figure S8: Ig-like domain mapping of the isolated antibodies.** Flow cytometry analysis of the binding of our new six full-length IgG2a reformatted antibodies to cells transiently transfected with mVCAM-1-eGFP WT (with or without N-terminal FLAG) or FLAG-tagged Ig-like deletion constructs. The secondary antibody only treatment is shown as negative control (Dylight 650). Displayed is the average normalized MFI-Max-Min (mVCAM-1-eGFP-positive, antibody positive cells normalized to MFI of the mVCAM-1-eGFP-negative, antibody negative cells) from one experiment with 2 replicates.

### Supplementary Tables

**Supplementary Table S1. Non-exhaustive list of antibodies developed against VCAM-1, their targeted domain (if known), and the associated function.** This table is only a snapshot of some of the reagents that have been developed to study VCAM-1 and it is not comprehensive. We apologise in advance as there are many antibodies and/or their functional assays that we have not included below.

| **Antibody** | **Domain targeted** | **Impact of antibody on VCAM-1 function** (domain associated) | **Reference** |
|---|---|---|---|
| 4B9 | 1 (And 4?)[1] | Inhibits adherence of PBL and lymphocytic cell line (not neutrophils) to activated HUVE. May be important for lymphocyte emigration and immune response. | [10-13] |
| ED11, GH12, and GE4 | 4, different epitopes | ED11 and GH12 can block VLA-4 dependent binding to VCAM-1 in certain genetic scenarios (e.g. absence of domain 1), although ED11 significantly blocked binding of Ramos cells to activated endothelial cells (ECs). | [12] |
| H6 | 1-2 | Blocks adhesion and transmigration between VCAM-1 and inflammatory cells. | [14] |
| 7H | 1-2 | Blocks adhesion and transmigration.<br>In ApoE$^{-/-}$ mice, 7H reduces the effect of atherosclerosis. | [14] |
| MVCAM.A[2] 429 | 1-2 | Reduces eosinophil infiltration on skin. | [15, 16] and [2] |
| M-/K1 and M-/K2[3] | 1 and 4 | Inhibit binding of VLA-4 expressing cells to VCAM-1.<br>Prevent lymphopoiesis.<br>Prevent binding of lymphoid cells to activated EC.<br>Reduces inflammation.<br>Enhances survival rates of grafts.<br>Blocks cardiomyocyte hypertrophy, myofibroblast activation, and cardiac remodelling (M/K2.7). | [17-21] and [3] |
| 2G7 | 1-3[4] | Blocks binding of T cells to activated ECs.<br>Blocks adhesion of basophils and eosinophils to activated ECs.<br>Blocks activation of CD4+ T cells. | [22-24] [5] |
| 1E7 | 1-3[4] | Does not block activation of CD4+ T cells.<br>Different epitope than 2G7. | [22, 24] [5] |
| 1G11, 1E5, 1.4C3, 6D9 | ? | 1G11 and 1E5 block T cell adhesion to ECs, but 14C3 and 6D9 do not.<br>All antibodies react with the circulatory VCAM-1 form, so they are not D4 only binders.<br>IG11 also blocks VCAM-1 mediated survival of neutrophils, and blocks VLA4-mediated release of interleukins by monocytes. | [25-27] |
| BBIG-V1[6] | 1 | Blocks VLA4-mediated release of interleukins by monocytes. | [27] |
| BBIG-V4 | 3 | | [28] |
| 19C3 | 2 | | [28, 29] |
| 1E10[7] | 1 | Likely BBIG-V3 | [28, 29] |
| E1/6 | 1 | Blocks binding to VLA4.<br>Prevents melanoma and other tumors adhering to activated ECs.<br>Inhibits binding of lymphoid cells to activated ECs.<br>Epitope of E1/6 is different from 4B9 (see above). | [11, 30-32] |
| Hu8/4 | 4-7 | Does not inhibit binding of melanoma to activated ECs. | [30] |
| VCAM-1 D6 Fab | 6 | Blocks leukocyte transendothelial migration. | [33] |

| | | | |
|---|---|---|---|
| 1/9 | Likely 1 and/or 4 | Blocks eosinophils rolling *in vivo* (likely by blocking domains 1 and/or 4). | [34] |
| 51-10C9 | 1 | Blocks leukocyte adhesion.<br>Enhanced TNF-α–stimulated EC production of IL-8. | [33] |
| 1.G11B1[8] | | Decrease in eosinophil tethering to IL-4 stimulated HUVECs particularly at high shear stress.<br>Inhibits cell adhesion between B cells and FLS (fibroblast-like σψνοϖιοχψτεσ). | [35, 36] |
| HD101 | 1 and 2 | Inhibits cell adhesion.<br>Induces VCAM-1 internalization into the cytoplasm.<br>Anti-inflammatory and anti-asthma effects. | [37] |
| **Antibody against VLA-4** | | | |
| *Natalizumab*[9] | $\alpha 4\beta 1$ *integrin blocker* | *Blocks interaction with VCAM-1 domains 1 and 4. Treatment for multiple sclerosis and Chron's disease.* | |

[1] Vonderheide and Springer [11] suggest that 4B9 binds to both Ig-like domains 1 and 4, but Osborn et al [38] data shows that it only binds 1 and suggests that it can sterically hinder interaction with domain 4 due to its size/position or possibly by perturbing the conformation/structure of the molecule upon binding.
[2] https://www.bdbiosciences.com/en-us/products/reagents/flow-cytometry-reagents/research-reagents/single-color-antibodies-ruo/fitc-rat-anti-mouse-cd106.553332
[3] https://www.merckmillipore.com/AU/en/product/Anti-VCAM-1-Antibody-clone-M-K,MM_NF-CBL1300?ReferrerURL=https%3A%2F%2Fwww.google.com%2F
[4] But could potentially also recognise other Ig-like domains that were not tested.
[5] From W. Newman, Otsuka America Pharmaceutical.
[6] https://www.rndsystems.com/products/human-vcam-1-cd106-antibody-bbig-v1_bba5
[7] https://www.rndsystems.com/products/human-vcam-1-cd106-fluorescein-conjugated-antibody-bbig-v3-ie10_bba22
[8] https://www.bio-rad-antibodies.com/monoclonal/human-cd106-antibody-1-g11b1-mca907.html?f=purified
[9] https://go.drugbank.com/drugs/DB00108

**Supplementary Table S2.** Plasmids constructed and used in this study.

| **Plasmid** | **Alias** |
|---|---|
| pUCIDT_mVCAM-1 | LZCBI 17 |
| pEGFP-N1_mVCAM-1 | LZCBI 18 |
| pET28b-1A9 | LZCBI 41 |
| pET28b-2D3 | LZCBI 45 |
| pET28b-2D8 | LZCBI 46 |
| pET28b-2E2 | LZCBI 47 |
| pET28b-2E6 | LZCBI 48 |
| pET28b-3H4 | LZCBI 50 |
| pET28b-LPETG | LZCBI 53 |
| pET28b-2D3-LPETG | LZCBI 54 |
| pET28b-2D8-LPETG | LZCBI 55 |
| pET28b-2E6-LPETG | LZCBI 56 |
| pET28b-3C12-LPETG | LZCBI 57 |
| pET28b-3H4-LPETG | LZCBI 58 |
| pET28b-1A9-LPETG | LZCBI 87 |
| pET28b-1H3-LPETG | LZCBI 88 |
| pET28b-2B9-LPETG | LZCBI 89 |
| pET28b-2E2-LPETG | LZCBI 90 |
| NBF320 JW8_ExpMG2a_2D3_Heavy chain | LZCBI 106 |
| NBF320 JW8_ExpMG2a_2E2_Heavy chain | LZCBI 108 |
| NBF320 JW8_ExpMG2a_2E6_Heavy chain | LZCBI 109 |
| NBF320 JW8_ExpMG2a_1A9_Heavy chain | LZCBI 110 |
| NBF320 JW8_ExpMG2a_3H4_Heavy chain | LZCBI 111 |
| NBF320 JW8_ExpMG2a_2D8_Heavy chain | LZCBI 112 |
| NBF372 JW8_ExpMG2a_2D3_Light chain | LZCBI 113 |
| NBF372 JW8_ExpMG2a_2E2_Light chain | LZCBI 115 |
| NBF372 JW8_ExpMG2a_2E6_Light chain | LZCBI 116 |
| NBF321 JW8_ExpMG2a_1A9_Light chain | LZCBI 117 |
| NBF372 JW8_ExpMG2a_2D8_Light chain | LZCBI 118 |
| NBF372 JW8_ExpMG2a_3H4_Light chain | LZCBI 119 |
| pEGFP-N1_mVCAM-1(1-24)-FLAG-mVCAM1(25-793) | LZCBI 120 |
| pEGFP-N1_mVCAM-1(1-24)-FLAG-mVCAM1(25-793)_ΔIgG_like_1 | LZCBI 121 |

| | |
|---|---|
| pEGFP-N1_mVCAM-1(1-24)-FLAG-mVCAM1(25-793)_ΔIgG_like_1-2 | LZCBI 122 |
| pEGFP-N1_mVCAM-1(1-24)-FLAG-mVCAM1(25-793)_ΔIgG_like_1-3 | LZCBI 123 |
| pEGFP-N1_mVCAM-1(1-24)-FLAG-mVCAM1(25-793)_ΔIgG_like_4-7 | LZCBI 124 |
| pEGFP-N1_mVCAM-1(1-24)-FLAG-mVCAM1(25-793)_ΔIgG_like_5-7 | LZCBI 125 |
| pEGFP-N1_mVCAM-1(1-24)-FLAG-mVCAM1(25-793)_ΔIgG_like_6-7 | LZCBI 126 |
| pEGFP-N1_mVCAM-1(1-24)-FLAG-mVCAM1(25-793)_ΔIgG_like_7 | LZCBI 127 |
| pEGFP-N1_mVCAM-1(1-24)-FLAG-mVCAM1(25-793)_ΔIgG_like_3 | LZCBI 129 |
| pEGFP-N1_mVCAM-1(1-24)-FLAG-mVCAM1(25-793)_ΔIgG_like_5 | LZCBI 130 |

**Supplementary Table S3.** Primers used in this study.

| Name | Sequence |
|---|---|
| *mVCAM1-FLAG Fwd* | ACGACGACGACAAGTTCAAGATCGAGATCAGC |
| *mVCAM1-FLAG Rev* | CCTTGTAGTCGGCCTGGGAAACGGCGA |
| *mVCAM1-d7 Fwd* | TTGGACGTGAAGGGGAAAGAGC |
| *mVCAM1-d7 Rev* | AGACACTTGGATGATCAGTTCC |
| *mVCAM1-d6 Rev* | GGCCACGTTCACGTACAGAGG |
| *mVCAM1-d5 Rev* | AGGGTCCTCAGGGAAGCTGTACAC |
| *mVCAM1-d4 Rev* | CTTTTCCTGCACGACCAGTTCCAC |
| *mVCAM1-d1 Fwd* | ATCTACTCATTTCCCAAGGACCCTG |
| *mVCAM1-d1 Rev* | CTTGTCGTCGTCGTCCTTGTAGTC |
| *mVCAM1-d2 Fwd* | ACAGTGAAGGAGCTCCAAGTG |
| *mVCAM1-d3 Fwd* | GAAAAGCCCTTCATCGTGG |
| *mVCAM1-d3 Rev* | GGAGATGTACACTTGGAGCTC |
| *mVCAM1-d5 Fwd* | GTGAACGTGGCCCCTAAGG |
| *LPETG codon corrected Fwd* | ACTGCCAGAAACCGGTCTCG |
| *LPETG codon corrected Rev* | GCGGCCGCAAGCTTGATATCC |

**Supplementary Table S4.** Protein parameters for purified scFv-LPETG antibodies.

| Sample | Molecular weight (Da) | Extinction Coefficient | Theoretical pI |
|---|---|---|---|
| 1A9 | 28038.20 | 52620 | 8.3 |
| 2D3 | 27980.87 | 57090 | 6.99 |
| 2D8 | 27337.19 | 46090 | 7.07 |
| 2E2 | 27183.86 | 43110 | 8.30 |
| 2E6 | 27976.79 | 54110 | 6.14 |
| 3H4 | 27209.86 | 54110 | 6.58 |

**Supplementary Table S5.** Summary of co-eluting proteins identified by Mass Spectrometry proteomics with Proteome Discoverer Software. The three proteins with the highest number of peptide spectrum matches (PSMs) are shown.

| **Accession #** | **Description** | **Coverage [%]** | **# Peptides** | **# PSMs** | **# Unique Peptides** | **# AAs** | **MW [kDa]** | **calc. pI** | **# Peptides (by Search Engine): Sequest HT** |
|---|---|---|---|---|---|---|---|---|---|
| P35340 | Alkyl hydroperoxide reductase subunit F OS=Escherichia coli (strain K12) OX=83333 GN=ahpF PE=1 SV=2 | 53 | 32 | 355 | 32 | 521 | 56.1 | 5.68 | 32 |
| P0A6Y8 | Chaperone protein DnaK OS=Escherichia coli (strain K12) OX=83333 GN=dnaK PE=1 SV=2 | 63 | 35 | 150 | 35 | 638 | 69.1 | 4.97 | 35 |
| P0A6F5 | 60 kDa chaperonin OS=Escherichia coli (strain K12) OX=83333 GN=groL PE=1 SV=2 | 57 | 28 | 129 | 28 | 548 | 57.3 | 4.94 | 28 |

We performed mass spectrometry proteomic analysis on one of the purified scFv samples to better understand the composition of the co-eluting proteins in the samples. The results showed three overly represented proteins (as determined by the number of peptide spectrum matches (PSMs) and % of coverage), shown in above. The number of PSMs identified positively correlates with the abundance of their associated protein in the sample. All three identified proteins have MW in the range of 55-70 kDa, which is where the strongest bands that co-elute essentially with all the purified scFvs and scFv-LPETGs appear (**Supplementary Figure S4A-C** and data not shown). Two of these three main proteins were chaperones, Chaperone protein DnaK and 60 kDa chaperonin, which help in protein folding. The third protein was an enzyme, Alkyl hydroperoxide reductase, that prevents DNA damage by alkyl hydroperoxides.
